## Supplemental files for "Feline Coronavirus Infection of Domestic Cats Causes Development of Cross-Reactive Antibodies to SARS-CoV-2 Receptor Binding Domain"

#### Supporting Information

**S1 Fig. The UF RBD and MassBiologic (MB) RBD sequences.** The information on the additional tags is shown with the complete RBD sequence (A). The MB-RBD is longer by 12 aa on the carboxyl-end. This 12 aa sequence is highly conserved among many human and animal coronaviruses, including FCoV and CCoV. The SCoV2 RBD has two strongly predicted O-glycosylation sites (bolded blue threonine **T** or serine **S**) and two N-glycosylation sites with low prediction (bold red asparagine **N**) on the amino-end on both UF and MB RBDs. MB-RBD has an additional O-glycosylation site on the first two aa residues (**S** and **T**) of the 12 aa addition on the carboxyl-end. The glycosylation prediction was based on a NetNGlyc 1.0 Server (https://services.healthtech.dtu.dk/service.php?NetNGlyc-1.0) for an N-glycosylation and NetOGlyc 4.1 Server (https://services.healthtech.dtu.dk/service.php?NetMHCpan-4.1) for O-glycosylation. Next, the sera from the same time-points as Figure 1E for the three toms were tested before the assay for Figures 1D/1E (B). Serum from SPF HOE serves as the negative control and serum from SPF cat DU1, which was vaccinated with FIV vaccine with residual BSA, serves as the anti-BSA Ab positive control.

**S2 Fig. The comparison of the aa residue numbers and aa sequence identity/similarity of the SCoV2 Wuhan (NC045512.1), FCoV1 UCD-1, and FCoV2 79-1146 (DQ010921.1) structural proteins.** The aa residue numbers are provided to estimate the MW of these proteins without glycosylation and to determine the aa sequence identity and similarity between the viruses (A). FCoV1 UCD-1 S (AB088222.1), NC (BAC1157.1) and M (BAC1156.1) were available but its Env sequence was unavailable, and therefore FCoV1 Dutch Cat1 Env (KX722530.1) was used in its place. The total aa identity or similarity value divided by the total aa residues including gaps x100 provides the percentage. The two values for S1 and S2 are provided based on where the SCoV2 S is processed into S1 and S2 at S1/S2 cleavage site and where the FCoV1 S and FCoV2 S are processed into S1 and S2 [44,45]. Under the % aa identity and % aa similarity for S1 and S2 proteins, the first value is for SCoV2 followed by the value for FCoV1 in row-1 and FCoV2 in row-2. The first value is for FCoV1 followed by the value for FCoV2 in row-3. The MW without glycan (B) is determined by using Peptide Analyzing Tool (https://www.thermofisher.com/us/en/home/life-science/protein-biology/peptides-proteins/custom-peptide-synthesis-services/peptide-analyzing-tool.html). The predicted N- and O-glycosylation numbers are determined by NetNGlyc 1.0 server and NetOGlyc 4.1 Server, respectively (B), and their network addresses are first described in Figure S1 legend.

**S3 Fig. Sequence alignment of SCoV2 Wuhan and FCoV2 79-1146 spike glycoproteins.** Clustal Omega 1.2.1 of JustBio Server (https://justbio.com/) was used for all sequence alignment analyses (A). Signal peptides for all sequences were determined by SignalP – 6.0 server of DTU Health Tech (https://services.healthtech.dtu.dk/service.php?SignalP). The abbreviations used include: amino-terminal domain (NTD), receptor binding domain (RBD), receptor binding motif (RBM), carboxyl-terminal domain (CTD), spike glycoprotein-1 (S1), and spike glycoprotein-2 (S2). RBM (underlined bold) has the aa residues that contact the cell receptor (e.g., species-specific ACE2 and APN). S1/S2 cleavage sites are found with cleavage motif (bold red) at the usual site for SCoV2 but the cleavage motif at the usual site is not found for FCoV2 (46,47). The S1/S2 cleavage site for FCoV2 and CCoV2 is reported to be amino-end adjacent to the fusion peptide, which is the S2' cleavage site for other strains. The FCoV2 79-1146 sequence is shown in magenta. Identical and similar aa residues are shown with an asterisk (*) for complete identity, strong similarity with a colon (:), and modest similarity with a single dot (.). The gaps are shown with a dash (-). The summary table (B) shows the total number of aa with gaps, number of aa with identity, number of aa with similarity, % aa identity, and % aa similarity. Two values for CTD, S1, and S2 are shown in the summary table (B) due to the major difference in the location of the S1/S2 cleavage site. In addition, the values for the proposed RBD sequence are also shown. The bolded % aa sequence similarities are the two lowest values and the two highest values.

**S4 Fig. Sequence alignment of SCoV2 Wuhan and FCoV2 UCD1 spike glycoproteins.** The sequences were aligned using the Clustal Omega 1.2.1 of JustBio Server (A) as described in Figure S2. The aa identity, similarity, and gap, and the abbreviations, cleavage site format, and table summary format are the same as those described in Figure S2. The FCoV1 UCD-1 sequence is shown in magenta. Two values for CTD, S1, and S2 are shown in the summary table (B) due to the major difference in the location of the S1/S2 cleavage site for these viruses. In addition, the values for the proposed RBD sequence are also shown. The bolded % aa sequence similarities are the two lowest values and the two highest values.

**S5 Fig. Anti-CoV analyses of group-housed laboratory and pet cats**. The analyses were performed using the FCoV2-WV and SCoV2 RBD ELISA (A) and the immunoblot strips (B,C) with SCoV2 UF-RBD (B1,C2 last row), proposed FCoV2 RBD (B2,C2 second row), FCoV2-WV (B3,C1), and the proposed FCoV1 RBD (B4,C2 first row). The eight group-housed laboratory cats (A,B) were unrelated to our UGA queens and UF toms, but the animal vendor was the same as the UGA queens and had different breeding lineages. Additionally, three group-housed pet cats from a household of five cats were similarly tested as preliminary study (C1-C4) to determine whether FCoV-infected pet cats develop cross-reactive antibodies to SCoV2 RBD. Pet cat sera KY1 and KY2 are from the same cat collected at the same time (KY1) as other two cats (KM1 and KN1) and 2.5 months later (KY2). The owner indicated that she has not been infected with SCoV2 and had no signs of COVID-19 (IRB202002902). However, her pet cats are allowed to roam outdoors, and therefore there is a remote possibility that they have been exposed to SCoV2 infected cat(s) outdoors. We have not tested her cats by SCoV2 RT-PCR or SCoV2 NAb assay. The serum from UGAQ4 was the $FCoV1^{+}$ control for FCoV2-WV (A,B3,C1) and FCoV1 RBD (B4,C2 first row) ELISA or immunoblot. The plasma or serum from cats J2a and UGA4.4 were $FCoV2^{+}$ (B2,C2 second row) and $SCoV2^{+}$ (B1,C2 last row) controls, respectively. The serum from cat UGAQ4 and UGA1.4 also served as weak positive controls for FCoV1 and FCoV2 RBD immunoblots respectively (C2 rows 1 & 2, B2 row 1), whereas serum from SPF cats HOF and HOE served as a negative control from all immunoblots and ELISA, respectively. All immunoblot photographs were adjusted to 10% brightness and 5% contrast for consistency.

**S6 Fig. Sequence alignment of SCoV2 Wuhan and SCoV1 Tor2 spike glycoproteins.** The sequences were aligned using the Clustal Omega 1.2.1 of JustBio Server as described in Figure S2 (A). The aa identity, similarity, and gap, and the abbreviations, cleavage site format, and table summary format are the same as those described in Figure S2. The RBD for each virus is shown in bold blue aa residues and the underlined section represents the RBM (A). These two sequences are highly conserved with a minimal number of gaps observed predominantly in NTD of S1 glycoprotein. The demarcation sites of the NTD, RBD, and CTD are identical between these viruses. The summary table (B) shows an additional column for the whole spike sequence. The bolded % aa sequence similarities are the one lowest value and the one highest value.

**S7 Fig. The SCoV2 UF2-RBD sequence with aa extensions.** The aa sequence extensions in blue aa residues were selected based on the sequence with aa conservation between SCoV2 and FCoV2 (S3 Fig) and SCoV2 Wuhan and SCoV1 Tor2 (S6 Fig) (A). The extensions were also based on the number of SCoV2-specific cytotoxic T lymphocyte (CTL) and T-helper ($T_H$) epitopes increased by the aa extensions as shown in S1 Table. Additionally, the aa residue number of UF2-RBD (269 aa) was increase to the residue number similar to our FCoV2 RBD (268 aa) that blocked *in vitro* FCoV2 infection more potently than the short SCoV2 UF-RBD. Our goal is to increase the CTL epitopes, which will eradicate the SCoV2 infected cells, but without increasing the $T_H$ epitopes, which has the potential to elevate the inflammatory responses. In our preliminary study (B), the sera from COVID-19 vaccinated humans (Y3, Y6) with no prior SCoV2 exposure were reacted with either SCoV2 UF2-RBD or UF-RBD immunoblot strip. Subject Y3 received three vaccinations with Pfizer S mRNA vaccine, while subject HY6 received first two vaccinations with Astra-Zeneca ChAsOX1-S recombinant vaccine and the last vaccination with Pfizer S mRNA vaccine. The year-2017 serum from Y3 was used as a negative human control. The sera from SCoV2-inoculated cat UGA4.1 and SPF cat HOF also served as positive and negative cat controls, respectively.

**Fig S8. Multi-sequence alignment of 10 CCoV2 spike *versus* four FCoV2 spike sequences.** The sequences were aligned using the Clustal Omega 1.2.1 of JustBio Server (A) as described in Figure S2. The aa identity, similarity, and gap, and the abbreviations, cleavage site format, and table summary format are the same as those described in Figure S2. The FCoV2 79-1146 sequence is shown in magenta. The RBD for FCoV2 79-1146 is shown with yellow highlight. Two values for CTD, S1, and S2 are shown in the summary table (B) due to the slight difference in the S1/S2 cleavage site motif among the CCoV2 and FCoV2 strains and are depicted with the cleavage motif. The bolded % aa sequence similarities are the one lowest value and the two highest values.

**Fig S9. Multi-sequence alignment of two CCoV1 spike *versus* 10 FCoV1 spike sequences.** The sequences were aligned using the Clustal Omega 1.2.1 of JustBio Server (A) as described in Figure S2. The aa identity, similarity, and gap, and the abbreviations, cleavage site format, and table summary format are the same as those described in Figure S2. The FCoV1 UCD-1 sequence is shown in magenta. The RBD for FCoV1 UCD-1 is shown with yellow highlight. Two values for CTD, S1, and S2 are shown in the summary table (B) due to the two aa residues difference in the S1/S2 cleavage site of the CCoV1 and FCoV1 and are designated by the virus. The bolded % aa sequence similarities are the one lowest value and the two highest values.

**S1 Table. CD8$^+$ CTL and CD4$^+$ T$_H$ Epitopes on SCoV2 UF-RBD and UF2-RBD.**

**S1 Fig**

**A** **MB-RBD (MassBiologics Wuhan RBD residue 319 to 541) glycoprotein kindly provided by MassBiologics with attached Tags:** c-Myc Peptide; 6xHis Tag

**UF-RBD from a plasmid with Wuhan RBD residue 319-529 sequence (MN975262.1) with attached Tags:** HRV 3C Cleavage Site; 8xHis Tag; Streptavidin-Binding Peptide (SBP) Tag

```
MB-RBD     319-RVQPTESIVRFPNITNLCPFGEVFNATRFASVYAWNRKRISNCVADYSVLYNSASFSTFK-378
UF-RBD     319-RVQPTESIVRFPNITNLCPFGEVFNATRFASVYAWNRKRISNCVADYSVLYNSASFSTFK-378
               ************************************************************

MB-RBD     379-CYGVSPTKLNDLCFTNVYADSFVIRGDEVRQIAPGQTGKIADYNYKLPDDFTGCVIAWNS-438
UF-RBD     379-CYGVSPTKLNDLCFTNVYADSFVIRGDEVRQIAPGQTGKIADYNYKLPDDFTGCVIAWNS-438
               ************************************************************

MB-RBD     439-NNLDSKVGGNYNYLYRLFRKSNLKPFERDISTEIYQAGSTPCNGVEGFNCYFPLQSYGFQ-498
UF-RBD     439-NNLDSKVGGNYNYLYRLFRKSNLKPFERDISTEIYQAGSTPCNGVEGFNCYFPLQSYGFQ-498
               ************************************************************

MB-RBD     499-PTNGVGYQPYRVVVLSFELLHAPATVCGPKKSTNLVKNKCVNF – c-Myc Peptide - 6xHis Tag
UF-RBD     499-PTNGVGYQPYRVVVLSFELLHAPATVCGPKK- HRV 3C Cleavage Site - 8xHis Tag - SBP Tag
               *******************************
```

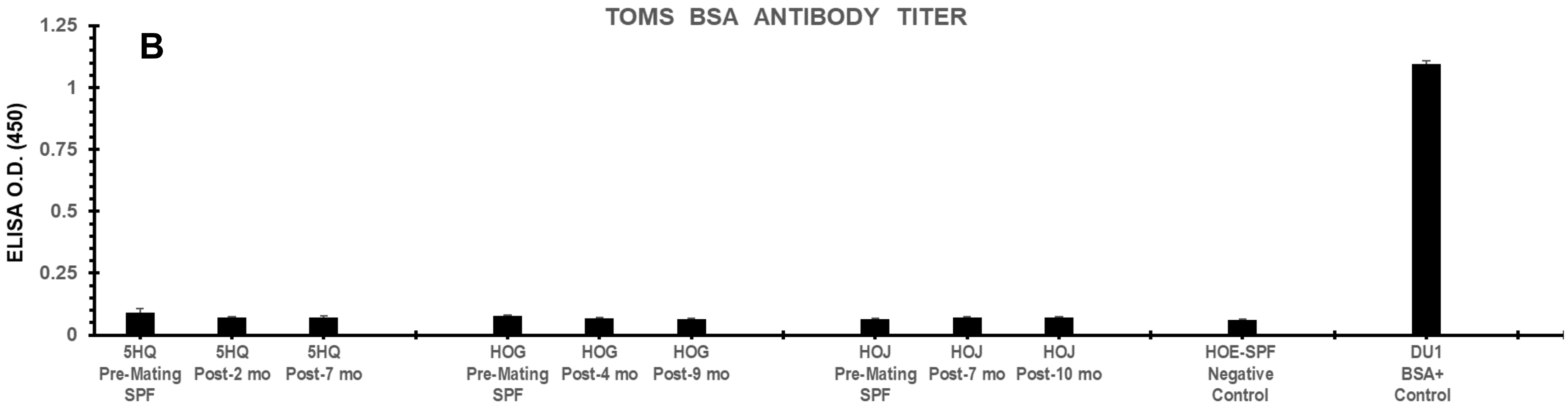

#### S2 Fig

##### A Summary of SCoV2 *versus* FCoV1/FCoV2 Structural Proteins

| SCoV2 *vs.* FCoV<br>FCoV1 *vs.* FCoV2 | Structural Protein * | Number of AA | AA Identity (%) | AA Similarity (%) |
|---|---|---|---|---|
| SCoV2 *vs.* FCoV1 | Spike (S1+S2) | 1273 *vs.* 1457 | 20.7% | 50.1% |
| SCoV2 *vs.* FCoV2 | Spike (S1+S2) | 1273 *vs.* 1452 | 22.0% | 50.4% |
| FCoV1 *vs.* FCoV2 | Spike (S1+S2) | 1457 *vs.* 1452 | 43.7% | 72.6% |
| SCoV2 *vs.* FCoV1 | S1 | 685 *vs.* 790 | 12.9% / 13.0% | 39.4% / 40.0% |
| SCoV2 *vs.* FCoV2 | S1 | 685 *vs.* 959 | 14.5% / 16.3% | 41.4% / 43.4% |
| FCoV1 *vs.* FCoV2 | S1 | 790 *vs.* 959 | 29.5% / 31.6% | 62.9% / 64.5% |
| SCoV2 *vs.* FCoV1 | S2 | 588 *vs.* 667 | 31.6% / 30.1% | 65.1% / 62.2% |
| SCoV2 *vs.* FCoV2 | S2 | 588 *vs.* 493 | 32.0% / 33.5% | 62.2% / 64.5% |
| FCoV1 *vs.* FCoV2 | S2 | 667 *vs.* 493 | 60.6% / 68.7% | 84.0% / 89.9% |
| SCoV2 *vs.* FCoV1 | Env | 75 *vs.* 82 | 25.6% | 68.3% |
| SCoV2 *vs.* FCoV2 | Env | 75 *vs.* 82 | 23.2% | 67.1% |
| FCoV1 *vs.* FCoV2 | Env | 82 *vs.* 82 | 96.3% | 97.6% |
| SCoV2 *vs.* FCoV1 | NC | 419 *vs.* 377 | 25.3% | 59.2% |
| SCoV2 *vs.* FCoV2 | NC | 419 *vs.* 377 | 25.7% | 58.0% |
| FCoV1 *vs.* FCoV2 | NC | 377 *vs.* 377 | 92.8% | 97.6% |
| SCoV2 *vs.* FCoV1 | M | 222 *vs.* 262 | 26.3% | 55.0% |
| SCoV2 *vs.* FCoV2 | M | 222 *vs.* 262 | 25.6% | 54.2% |
| FCoV1 *vs.* FCoV2 | M | 262 *vs.* 262 | 95.4% | 98.5% |

* Spike 1 (S1), Spike 2 (S2), Envelope (Env), Nucleocapsid (NC) and Membrane (M) proteins.

##### B Structural Proteins Based on Total AA Number and the Predicted N- and O-Glycosylation(s)

| Structural Protein | FCoV1 (Da)* | FCoV1 Glycan N / O† | FCoV2 (Da)* | FCoV2 Glycan N / O† |
|---|---|---|---|---|
| S (S1+S2) | 163,727 | 3 / 3 | 160,473 | 4 / 1 |
| S1 | 89,666 | 3 / 1 | 106,185 | 4 / 1 |
| S2 | 74,080 | 1 / 2 | 54,307‡ | 1 / 0 |
| Env | 9,379 | 0 / 0 | 9,371 | 0 / 0 |
| NC | 42,663 | 0 / 0 | 42,703 | 0 / 4 |
| M | 29,920 | 1 / 0 | 29,832 | 1 / 0 |

* The MW of FCoV is determined by the total aa residue number using the Peptide Analyzing Tool server.

† The number of N-glycosylation (N) with high or medium potential and O-glycosylation (O) with strong potential are shown as (N / O).

‡ The predicted FCoV2 S2 shown in Fig. 2*B* matches the MW of FCoV1 S2 with glycan(s). Hence, the predicted FCoV2 S1/S2 cleavage site may be more closer to that of FCoV1, and FCoV2 S2 with glycan may be closer in size to the FCoV1 S2 with glycan(s).

### S3 Fig. SCoV2 Wuhan *versus* FCoV2 WSU79-1146 Spike Glycoproteins

## A

```
                                  Signal Peptide         NTD / S1
YP009724390.1_SCoV2_Wuhan         MFVF----------LVLLPLVSSQCVNLTTRT-----QLPP-------AYTNSFTRGVYY
AAY32596.1_FCoV2_WSU79-1146       MIVLVTCLLLLCSYHTVLSTTNNECIQVNVTQLAGNENLIRDFLFSNFKEEGSVVVGGYY
                                  *:*:           .:*  ...:*:::..        :*              *.. * **
                                                       NTD

YP009724390.1_SCoV2_Wuhan         PDKVFRSSVLHSTQDLFLPFFSNVTWFHA-IH-VSGTNGTKRFDNPVLPFNDGV------
AAY32596.1_FCoV2_WSU79-1146       PTEVWYNCSRTARTTAF-QYFNNIHAFYFVMEAMENSTGNAR-GKPLLFHVHGEPVSVII
                                  * :*: ..   :    *  :*.*:  *:  :. :. :.*. *  :*:*  . .*

YP009724390.1_SCoV2_Wuhan         --YFASTEKSNII--------------------RGWIFGTTLDSKTQSL--LIVNNATNV
AAY32596.1_FCoV2_WSU79-1146       SAYRDDVQQRPLLKHGLVCITKNRHINYEQFTSNQWNSTCTGADRKIPFSVIPTDNGTKI
                                    *  ..::  ::                    . *    *  .:.  :  : .:*.*::

YP009724390.1_SCoV2_Wuhan         VIKVCEFQFCNDPFLGVYYH-KNNKSWMESEFRVYSSANNCTFEYVSQPFLMDLEG----
AAY32596.1_FCoV2_WSU79-1146       YGLEWNDDFVTAYISGRSYHLNINTNWFNNVTLLYSRSSTATWEYSAAYAYQGVSNFTYY
                                       : :*  .  : *  ** : *..*::.  :** :...*:** :      :.

YP009724390.1_SCoV2_Wuhan         KQGNFKNLREFV----FKNIDGYFKIYSKHTPINLVRDLPQGFSALEPLVDLP-IGINIT
AAY32596.1_FCoV2_WSU79-1146       KLNNTNGLKTYELCEDYEHCTGYATNVFAP---TSGGYIPDGFSFNNWFLLTNSSTFVSG
                                  *  * : *: :     ::.  ** .         .    :*:***  : ::      :

YP009724390.1_SCoV2_Wuhan         RFQTLLALHRSYLTPG----DSSSGWTA-----------------GAAAYYVGYL---Q-
AAY32596.1_FCoV2_WSU79-1146       RFVTNQPLLINCLWPVPSFGVAAQEFCFEGAQFSQCNGVSLNNTVDVIRFNLNFTADVQS
                                  ** *    *  . * *       ::. :                 .  : : :    *

                                                                     NTD       RBD / S1
YP009724390.1_SCoV2_Wuhan         -PRTFLLKYNENGTITDAVDCALDPLSETKCTLKSFTVEKGIYQTSNFRVQPTESIVRFP
AAY32596.1_FCoV2_WSU79-1146       GMGATVFSLNTTGGVILEISCYSDTVSESSSYSY-GEIPFGITDGPRY------CYVLY-
                                     : ::. * .* :   :.*  * :**:..    :  ** :  .:      . * :
                                                                   NTD

YP009724390.1_SCoV2_Wuhan         NITNLCPFGEVFNATRFASVYAWNRKRIS----------NCVA-----------------
AAY32596.1_FCoV2_WSU79-1146       NGTALKYLGTLPPSVKEIAISKWGHFYINGYNFFSTFPIGCISFNLTTGVSGAFWTIAYT
                                  * * *   :* :   :.:  ::  * :  *.            *::
                                  Proposed RBD / S1

YP009724390.1_SCoV2_Wuhan         -------------DYSVLYNSASFSTFKCYGVSPT--------KLNDLCFTN---VYADS
AAY32596.1_FCoV2_WSU79-1146       SYTEALVQVENTAIKNVTYCNSHINNIKCSQLTANLNNGFYPVASSEVGFVNKSVVLLPS
                                               .* * .: :..:** :: .           .:: *.*    *    *

YP009724390.1_SCoV2_Wuhan         FVIRGDEVRQIAPGQTGKIADYNYKLPDDFTGCVIAWNSNNLDSKVGGNYNYLYRLFRKS
AAY32596.1_FCoV2_WSU79-1146       FFTYTA--VNITIDLGMKLSGYGQPIASTLSNITLPMQDNNTDVYCIRS--NQFSVYVHS
                                  *.         :*:      *:: *    : . ::  .:  :.** *      .   : :: :*

YP009724390.1_SCoV2_Wuhan         NLK------PFERDI-----STEIYQAGSTPC---------------N--GVEGFNCYFP
AAY32596.1_FCoV2_WSU79-1146       TCKSSLWDNIFNQDCTDVLEATAVIKTGTCPFSFDKLNNYLTFNKFCLSLSPVGANCKFD
                                  . *         *::*        :* : ::*: *                  .  * ** *

                                                                        RBD            CTD / S1
YP009724390.1_SCoV2_Wuhan         LQSYGFQPTNGVGYQPYRVVVLSFELLHAPATVCGPKKSTNLVKNKCVNFNFNGLTGTGV
AAY32596.1_FCoV2_WSU79-1146       VAARTRT-NEQVVRSLYVIYEEGDNIVGVPSDNSGLHDLSVLHLDSCTDYNIYGRTGVGI
                                  : :      .: *   . * :    . ::: .*: .* :. : *   :.*.:: *: * **.*:
                                                              Proposed RBD          CTD

YP009724390.1_SCoV2_Wuhan         LTESNKKFLPFQQFGRDIADTTDAVRDPQTLEILDITPCSFGGVSVITPGTNTSNQVAVL
AAY32596.1_FCoV2_WSU79-1146       IRRTNSTLLSGLYYTSLSG-DLLGFKNVSDGVIYSVTPCDVSVQAAVIDGAIVGAMTS--
                                  : .:*..:*     :      .    ..:: .     *  .:***...  :.:   *: ..  .:

YP009724390.1_SCoV2_Wuhan         YQDVNCTEVPVAIHADQLTPTWRVYSTGSNVFQTRAGCLIGAEHVNNSYECDIPIG---A
AAY32596.1_FCoV2_WSU79-1146       ---INSE--LLGLTHWTTTPNFYYYSIYNYTSERTRGTAI----DSNDVDCEPVITYSNI
                                     :*.     :.:      **.:  **  . . :    *  *      .*. :*:  *

                                         CTD
                                       ← S1/S2 →
YP009724390.1_SCoV2_Wuhan         GICASYQTQTNSPRRARSVASQSIIAYTMSLGAENSVAYSNNSIAIPTNFTISVTTEILP
AAY32596.1_FCoV2_WSU79-1146       GVCKN----------------GALVFINVTHSDGDVQPISTGNVTIPTNFTISVQVEYMQ
                                  *:* .                     :::  .:: .  :     *. .::********* .* :
```

```
YP009724390.1_SCoV2_Wuhan        VSMTKTSVDCTMYICGDSTECSNLLLQYGSFCTQLNRALTGIAVEQDKNTQE-VF-----
AAY32596.1_FCoV2_WSU79-1146      VYTTPVSIDCARYVCNGNPRCNKLLTQYVSACQTIEQALAMGARLENMEVDSMLFVSENA
                                 *  * .*:**: *:*  . .*.:** ** * *  :::**:  *  :: :.:. :*

                                                                                  S2'
YP009724390.1_SCoV2_Wuhan        ------------AQVKQIYKTPPIKDFGGF---NFSQILPD-PSKPSKRSFIEDLLFNKV
AAY32596.1_FCoV2_WSU79-1146      LKLASVEAFNSTENLDPIYKEWPS--IGGSWLGGLKDILPSHNSKRKYGSAIEDLLFDKV
                                             ::. ***  *   :**     ..:***.  ** .  * ******:**
                                                                         S1/S2
                                                                    CTD

YP009724390.1_SCoV2_Wuhan        TLADAGFI-KQYGDCLGDIAARDLICAQKFNGLTVLPPLLTDEMIAQYTSALLAGTITSG
AAY32596.1_FCoV2_WSU79-1146      VTSGLGTVDEDYKRCTGGYDIADLVCAQYYNGIMVLPGVANADKMTMYTASLAGGI-TLG
                                 . :   * : ::*  * *     **:*** :**: *** :  . : :: **::* .*  * *

YP009724390.1_SCoV2_Wuhan        WTFGAGAALQIPFAMQMAYRFNGIGVTQNVLYENQKLIANQFNSAIGKIQDSL-------
AAY32596.1_FCoV2_WSU79-1146      A--LGGGAVAIPFAVAVQARLNYVALQTDVLNKNQQILANAFNQAIGNITQAFGKVNDAI
                                     .*.*: ****: :   *:* :.:   :** :**:::** **.***:* :::

YP009724390.1_SCoV2_Wuhan        -------SSTASALGKLQDVVNQNAQALNTLVKQLSSNFGAISSVLNDILSRLDKVEAEV
AAY32596.1_FCoV2_WSU79-1146      HQTSQGLATVAKALAKVQDVVNTQGQALSHLTVQLQNNFQAISSSISDIYNRLDELSADA
                                        ::.*.**.*:***** :.***. *. **..** **** :.** .***::.*:.

YP009724390.1_SCoV2_Wuhan        QIDRLITGRLQSLQTYVTQQLIRAAEIRASANLAATKMSECVLGQSKRVDFCGKGYHLMS
AAY32596.1_FCoV2_WSU79-1146      QVDRLITGRLTALNAFVSQTLTRQAEVRASRQLAKDKVNECVRSQSQRFGFCGNGTHLFS
                                 *:********  :*:::*:* * * **:***  :**   *:.***  .**:*.  ***:* **:*

YP009724390.1_SCoV2_Wuhan        FPQSAPHGVVFLHVTYVPAQEKNFTTAPAICHDGKA-HFP------REGVFVSNGTHWFV
AAY32596.1_FCoV2_WSU79-1146      LANAAPNGMIFFHTVLLPTAYETVTAWSGICASDGDRTFGLVVKDVQLTLFRNLDDKFYL
                                 : ::**.*::*:*.. :*:  :..*:  .** .       *         :  :* .   ::::

YP009724390.1_SCoV2_Wuhan        TQRNFYEPQIITTDNTFVSGNCDVVIGIVNNTVYDPLQPELDS----FKEELDKYFKNHT
AAY32596.1_FCoV2_WSU79-1146      TPRTMYQPRVATSSDFVQIEGCDVLFVNATVIDLPSIIPDYIDINQTVQDILENYEPNWT
                                 * *.:*:*:: *:.: .     ***::   ..       : *:  .    .:: *::*   * *

YP009724390.1_SCoV2_Wuhan        SPDVDLGDISGINASVVNIQKEI--------------DRLNEVAKNLNESLIDLQELGKY
AAY32596.1_FCoV2_WSU79-1146      VPEFTLDIF---NATYLNLTGEIDDLEFRSEKLHNTTVELAILIDNINNTLVNLEWLNRI
                                  *:. *   :    **: :*:   **                .*  : .*:*::*::*: * :

YP009724390.1_SCoV2_Wuhan        EQYIKWPWYIWLGFIAGLIAIVMVTIMLCCM-TSCCSCLK---GCCSCGSCCKFDEDDSE
AAY32596.1_FCoV2_WSU79-1146      ETYVKWPWYVWLLIGLVVVFC-IPLLLFCCFSTGCCGCIGCLGSCCH-SICSRRQFENYE
                                 * *:*****:** :   ::   :  :::**: *.**.*:    .**   . *.: : :: *

YP009724390.1_SCoV2_Wuhan        PVLKGVKLHYT
AAY32596.1_FCoV2_WSU79-1146      PI-EKVHVH--
                                 *: : *::*
```

**B**

**Summary of Amino Acid (AA) Sequence Identity and Similarity between SCoV2 and FCoV2**

| | **NTD** | **RBD** | **RBD (Proposed)** | **CTD (SCoV2)** | **CTD (FCoV2)** | **S1 (SCoV2)** | **S1 (FCoV2)** | **S2 (SCoV2)** | **S2 (FCoV2)** |
|---|---|---|---|---|---|---|---|---|---|
| Total No. of AA | 370 | 290 | 273 | 147 | 297 | 834 | 984 | 654 | 504 |
| No. of AA with Identity | 55 | 38 | 36 | 24 | 64 | 121 | 161 | 210 | 170 |
| No. of AA with Similarity | 155 | 107 | 100 | 69 | 151 | 345 | 427 | 409 | 326 |
| % Identity | 14.9% | 13.1% | 13.2% | 16.8% | 21.5% | 14.5% | 16.4% | 32.1% | 33.7% |
| % Similarity | 41.9% | **36.9%** | **36.6%** | 54.7% | 50.8% | 41.4% | 43.4% | **62.5%** | **64.7%** |

### S4 Fig. SCoV2 Wuhan *versus* FCoV1 UCD1 Spike Glycoproteins

## A

```
                                 Signal Peptide  NTD / S1
YP009724390.1_SCoV2_Wuhan        -MFVFLVLLPLVSSQCVNLTTRTQLPPAYTNSFTRGV----YYP-DKVFRS-SVLHSTQD
AB088222.1_FCoV1_UCD-1           MILLIFALFSVVRSEDA---PHGVTLPHFNTSYNNPKFELNFYNFLQTWDIPPNTETILG
                                  :::::.*:  :*  *:  .    :     *  :..*:..       :*    :.:       .:
                                               NTD

YP009724390.1_SCoV2_Wuhan        LFLPFF---SNVTWFHAIHVSGTNG----TKRFDNPVLPFNDGVYFASTEKSNIIRGWIF
AB088222.1_FCoV1_UCD-1           GYLPYCRTGQNCGWYNFVYNHAVQNAKYSYINTQNLNIPNVHGVYFDVREHNS-DGVWDT
                                  :**:     .*   *:. ::   ..:         .  :*   :*   .****    *:..     *

YP009724390.1_SCoV2_Wuhan        --------GTTLDSKTQSLL-----IVNNATNVVIKVCEFQFCNDP--------------
AB088222.1_FCoV1_UCD-1           RDRVGLLIAIHGESHYSLLMVLQDNVEENQPHVAVKICHWQLGNISSYHQFNLNLGDGGQ
                                         .   :*:  . *:      :  :*   .*.:*:*.:*: *

YP009724390.1_SCoV2_Wuhan        --------------------------FLGVYYH-KNNKSWMESEFRVYSSANNCTFEYVS
AB088222.1_FCoV1_UCD-1           CVFNQRFSLDTVLTANDFYGFQWTDTYVDIYLGGTITKVWVDNDWSVVEASISYHWSRLN
                                                           :: :*    .  .* *::.:: * .:: .   :. :.

YP009724390.1_SCoV2_Wuhan        QPFLMDLEGKQ----------GNFKNLREFVFKNIDGYFKIYSKHTPINLVRDLPQGFSA
AB088222.1_FCoV1_UCD-1           YGYYMQFVNRTTYYIYNNTGGSNYTHLH--LQECHSDYCAGYAKNVFVPIDGKIPESFSF
                                   : *::   :             .*:..*:  : :   .  *    *:*.. : :   .:*:.**

YP009724390.1_SCoV2_Wuhan        ---------------------------LEPLVDLPIGINITRFQTLLALHRSYLTPGDSS
AB088222.1_FCoV1_UCD-1           NNWFLLSDKSTLVQGRVLSSQPVFVQCLRPVPTLSNNTAVVHFN------NDVFCPNV--
                                                            *.*:   *       :.:*:       .. : *

                                                                                  NTD
YP009724390.1_SCoV2_Wuhan        SGWTAGAAAYYVGYLQPRT--------FLLKYNENGTITDAVDCALDPLSETKCTLKSFT
AB088222.1_FCoV1_UCD-1           ---TADVLRFNLNFSDTDVYTASTNDQLYFTFEDNT--TASIACYS--------------
                                    ** .  : : : :   .          : :.:::*    * :: *

                                               RBD / S1
YP009724390.1_SCoV2_Wuhan        VEKGIYQTSNFR-VQPTESIVRFPNITNLCPFGEVFNATRFASVYAWN--RK---RISNC
AB088222.1_FCoV1_UCD-1           -------SANVTDNQP------ANNSTSHIPFGKTMHSYFCFANFSNSMVRQFLGILPPT
                                        ::*.   **          * *.  ***:.:.:    : :: .   *:     :
                                                      NTD    Proposed RBD / S1

YP009724390.1_SCoV2_Wuhan        VADYSVLYN-SASFSTFKCYGVSPTKLNDLC-------------FTNVYADSFVIRGDEV
AB088222.1_FCoV1_UCD-1           VREFAFGRDGSIFVNGYKYFSLPPIKSVNFSIGSVEQHGFWTIAYTNYTDVMVDINGTGI
                                 * :::.  : *   .. :* :.: * *   ::.              :**      . *.*  :

YP009724390.1_SCoV2_Wuhan        RQI----APGQTGKIADYNYKLPDDFTGCV------------------IAWNSNNLDSKVG
AB088222.1_FCoV1_UCD-1           TRLFYCDSPINRIKCQQLKHELPDGFYAASMLVKKDLPKTFVTMPQFYNWMTVTLHVVLN
                                  ::    :* :   *   : :::*** *  ..                         * : .*.   :

YP009724390.1_SCoV2_Wuhan        GNYNYLYRLFRKSNLKPFERDISTEIYQAGS-----------------------------
AB088222.1_FCoV1_UCD-1           DTEKKADIILAKAPELASLADIHFEIVQANGSVANVTSLCVQTRQIALFYKYTSLQSLYT
                                  . :     :: *:         **   ** ** .

YP009724390.1_SCoV2_Wuhan        -TPCNGVEGFNCYFPLQS---------YGFQPTNG-----------VGYQ-PYRVVVLSF
AB088222.1_FCoV1_UCD-1           YSNLVELQNYDCPFSPQQFNNYLQFETLCLDTNPAVAGCKWALVHDVKWRTQFATITVSY
                                  :    :: ::* *  *.              :: . .              * ::  : .:.:*:

                                              RBD            CTD
YP009724390.1_SCoV2_Wuhan        ----ELLHAPATVCGPKKSTNLVKNKCVNFNFNGLTGTGVLTESNKKFLPFQQFGRDIAD
AB088222.1_FCoV1_UCD-1           KDGAMITTMPKAQLGFQDISNLVKDECTDYNIYGFHGTGIIRNTTSRLVAGLYY-TSISG
                                       :   * :   * :. :****::*.:: *: *: ***:: ::..:::     :  .*:
                          Proposed RBD                 CTD

YP009724390.1_SCoV2_Wuhan        TTDAVRDPQTLEILDITPCSFGGVSVITPGTNTSNQVAVLYQDVNCTEVPVAIHADQLTP
AB088222.1_FCoV1_UCD-1           DLLAFKNSTTGEIFTVVPCDLTAQAAVINDEIVG-----VITAVNQTDLFEFVNH-----
                                    *.::  * **: :.**.: . :.:       ..      :   ** *::    :.

                                                                                        CTD
                                                                                     S1/S2
YP009724390.1_SCoV2_Wuhan        TWRVYSTGSN-----VFQTRAGCLIGAEHVNNSYECDIPIGAGICASYQTQTNSPRRARS
AB088222.1_FCoV1_UCD-1           THSRRSRGSTSTSVTTYTMPQFYYITKWNNDTSTNCTSVITYSS--------------FA
                                 *     * **.     .:         *    . :.* :*    *  .                  :
                                      S1/S2
                                 CTD
```

```
YP009724390.1_SCoV2_Wuhan       VASQSIIAYTMSLGAENSVAYSNNSIAIPTNFTISVTTEILPVSMTKTSVDCTMYICGDS
AB088222.1_FCoV1_UCD-1          ICNTGEIKYVKDDSIGVIKPISTGNISIPKNFTVAVQAEYIQIQVKPVVLDCAKYVCNGN
                                :.. . * *. . .       *. .*:**.***::* :* : :.:. . :**: *:*  .

YP009724390.1_SCoV2_Wuhan       TECSNLLLQYGSFCTQLNRALTGIAVEQDKNTQE---------VFAQVKQI-YKTPPIKD
AB088222.1_FCoV1_UCD-1          SHCLSLLTQYTSACQTIENALNLGARLESLMLNDMITVSSRSLELATVERFNATAPGGEK
                                :.* .** ** * *  ::.**.  *  :.   ::          :* *:::  .:*  :.

                                                   S2'
YP009724390.1_SCoV2_Wuhan       FGGFNFSQIL-PDPSKPSKRSFIEDLLFNKVTLADAGFI-KQYGDCLGDIAARDLICAQK
AB088222.1_FCoV1_UCD-1          LGGLYFDGLSSLLPPRVGQRSAVEDLLFNKVVTSGLGTVDDDYKKCSAGTDVADLVCAQY
                                :**: *. :    * : .:** :********. :  * : .:* .* .   . **:***
                                                   S2'

YP009724390.1_SCoV2_Wuhan       FNGLTVLPPLLTDEMIAQYTSALLAGTITSGWTFGAGAALQIPFAMQMAYRFNGIGVTQN
AB088222.1_FCoV1_UCD-1          YNGIMVLPGVVDGNKMAMYTASLIGGMALGS----ITSAVAVPFAMQVQARLNYVALQTD
                                :**: *** ::  : :* **::*:.*   ..      :*: :*****:  *:* :.:  :

YP009724390.1_SCoV2_Wuhan       VLYENQKLIANQFNSAIGKIQD--------------SLSSTASALGKLQDVVNQNAQALN
AB088222.1_FCoV1_UCD-1          VLQENQKILANAFNNAIGNITLALGKVSNAITTISDGFNTMASALTKIQSVVNQQGEALS
                                ** ****::** **.***:*                .:.: **** *:*.****:.:**.

YP009724390.1_SCoV2_Wuhan       TLVKQLSSNFGAISSVLNDILSRLDKVEAEVQIDRLITGRLQSLQTYVTQQLIRAAEIRA
AB088222.1_FCoV1_UCD-1          LLTSQLRKNFQAISSSIAEIYNRLEKVEADAQVDRLITGRLAALNAYVSQTLTQYAEVKA
                                 *..** .** **** : :* .**:****:.*:******** :*::**:* * : **::*

YP009724390.1_SCoV2_Wuhan       SANLAATKMSECVLGQSKRVDFCGKGYHLMSFPQSAPHGVVFLHVTYVPAQEKNFTTAPA
AB088222.1_FCoV1_UCD-1          SRQLAMEKVNECVKSHSDRYGFCGDGTHLFSLVNSAPDGLLFFHTVLLPTEWEEVTAWSG
                                * :**  *:.*** .:*.*  ***.* **:*: :***.*::*:*.. :*:: ::.*:  .

YP009724390.1_SCoV2_Wuhan       ICHDGKAHFPREG---VFVSNGTHWFVTQRNFYEPQIITTDNTFVSGNCDVVIGIVNNTV
AB088222.1_FCoV1_UCD-1          ICVNDTFAYVLKDFEYFIFSYNNTYMVTPRNMFQPRKPQMSDFVQIMSCEVTFLNTTYTT
                                ** : .  :  :    .:.*  . ::** **:::*:    .: .   .*:*.:  ..  *.

YP009724390.1_SCoV2_Wuhan       YDPLQP----ELDSFKEELDKYFKNHTSPDVDLGDISGINASVVNIQKE-----------
AB088222.1_FCoV1_UCD-1          FQEIVIDYIDINKTIADMLEQYYSNYTTPELDLQLE-IFNQTKLNLTAEIDQLEQRADNL
                                :: :        .:: : *::*:.*:*:*::**     :* : :*:  *

YP009724390.1_SCoV2_Wuhan       ---IDRLNEVAKNLNESLIDLQELGKYEQYIKWPWYIWLGFIAGLIAIVMVTIMLCCM-T
AB088222.1_FCoV1_UCD-1          TNIAHELQEYIDNLNKTLVDLEWLNRIETYVKWPWYVWLLIGLVVV-FCIPLLLFCCLST
                                    ..*:*  .***::*:**: * : * *:*****:** :   :: : :  :::**: *

YP009724390.1_SCoV2_Wuhan       SCCSCLK---GCCSCGSCCKFDEDDSEPVLKGVKLHYT
AB088222.1_FCoV1_UCD-1          GCCGCFGCLGSCCH-SLCSRRQFENYEPI-EKVHIH--
                                .**.*:    .**  . *.: : :: **: : *::*
```

**B**

**Summary of Amino Acid (AA) Sequence Identity and Similarity between SCoV2 and FCoV1**

| | **NTD** | **RBD** | **RBD (Proposed)** | **CTD (SCoV2)** | **CTD (FCoV1)** | **S1 (SCoV2)** | **S1 (FCoV1)** | **S2 (SCoV2)** | **S2 (FCoV1)** |
|---|---|---|---|---|---|---|---|---|---|
| Total No. of AA | 402 | 307 | 279 | 149 | 97 | 885 | 833 | 639 | 691 |
| No. of AA with Identity | 46 | 39 | 33 | 23 | 17 | 108 | 114 | 203 | 209 |
| No. of AA with Similarity | 154 | 111 | 101 | 64 | 50 | 330 | 344 | 417 | 431 |
| % Identity | 11.4% | 12.7% | 11.8% | 15.4% | 17.5% | 12.2% | 13.7% | 31.8% | 30.2% |
| % Similarity | 38.3% | **36.2%** | **36.2%** | 42.9% | 51.5% | 37.3% | 41.3% | **65.3%** | **62.4%** |

S5 Fig

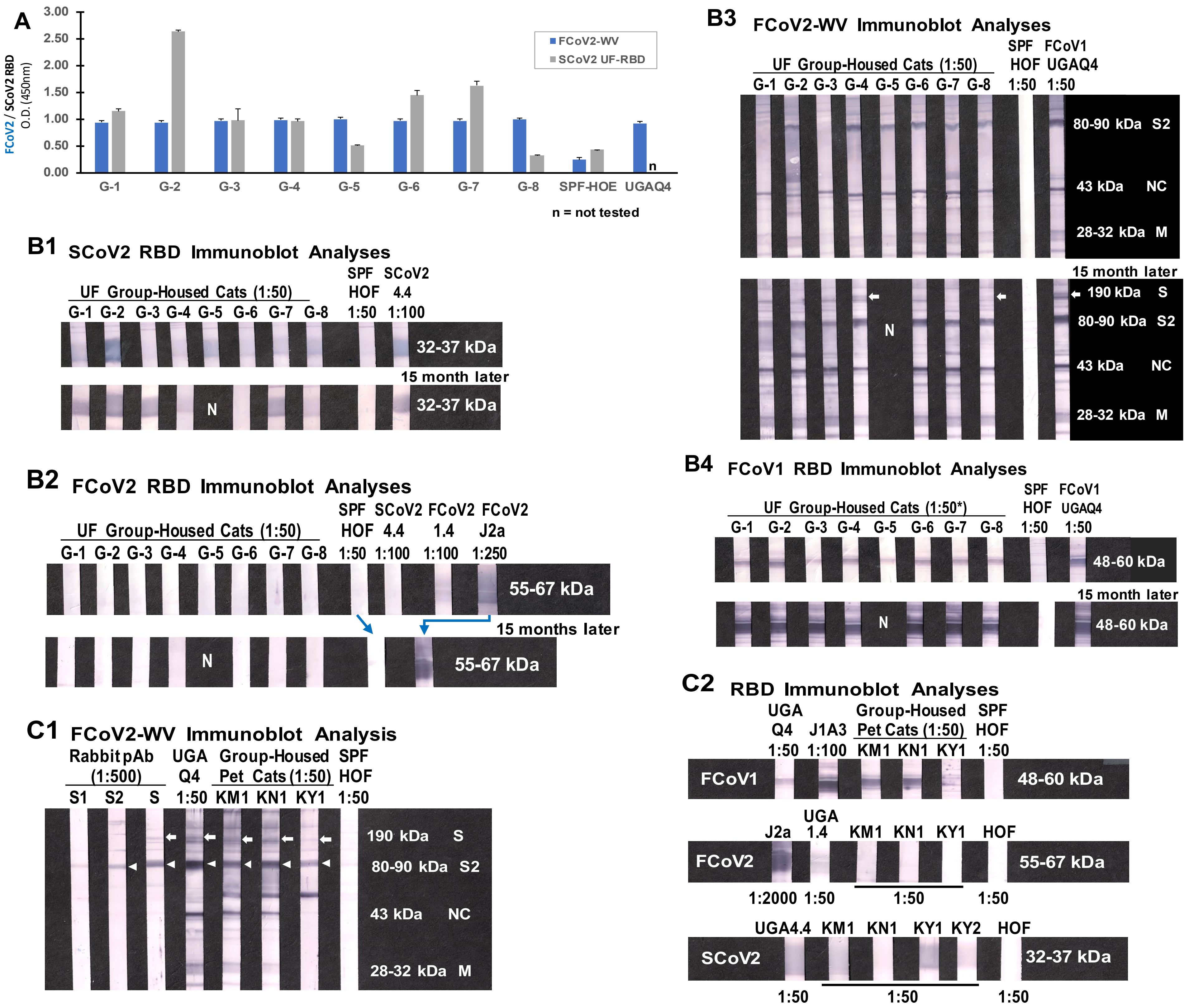

A
FCoV2 / SCoV2 RBD O.D. (450nm)
3.00
2.50
2.00
1.50
1.00
0.50
0.00
FCoV2-WV
SCoV2 UF-RBD
n
G-1 G-2 G-3 G-4 G-5 G-6 G-7 G-8 SPF-HOE UGAQ4
n = not tested
B1 SCoV2 RBD Immunoblot Analyses
UF Group-Housed Cats (1:50)
SPF HOF 1:50
SCoV2 4.4 1:100
G-1 G-2 G-3 G-4 G-5 G-6 G-7 G-8
32-37 kDa
15 month later
N
32-37 kDa
B2 FCoV2 RBD Immunoblot Analyses
UF Group-Housed Cats (1:50)
SPF HOF 1:50
SCoV2 4.4 1:100
FCoV2 1.4 1:100
FCoV2 J2a 1:250
G-1 G-2 G-3 G-4 G-5 G-6 G-7 G-8
55-67 kDa
15 months later
N
55-67 kDa
C1 FCoV2-WV Immunoblot Analysis
Rabbit pAb (1:500)
S1 S2 S
UGA Q4 1:50
Group-Housed Pet Cats (1:50)
KM1 KN1 KY1
SPF HOF 1:50
190 kDa S
80-90 kDa S2
43 kDa NC
28-32 kDa M
B3 FCoV2-WV Immunoblot Analyses
UF Group-Housed Cats (1:50)
G-1 G-2 G-3 G-4 G-5 G-6 G-7 G-8
SPF HOF 1:50
FCoV1 UGAQ4 1:50
80-90 kDa S2
43 kDa NC
28-32 kDa M
15 month later
N
190 kDa S
80-90 kDa S2
43 kDa NC
28-32 kDa M
B4 FCoV1 RBD Immunoblot Analyses
UF Group-Housed Cats (1:50*)
G-1 G-2 G-3 G-4 G-5 G-6 G-7 G-8
SPF HOF 1:50
FCoV1 UGAQ4 1:50
48-60 kDa
15 month later
N
48-60 kDa
C2 RBD Immunoblot Analyses
UGA Q4 1:50
J1A3 1:100
Group-Housed Pet Cats (1:50)
KM1 KN1 KY1
SPF HOF 1:50
FCoV1
48-60 kDa
J2a
UGA 1.4
KM1 KN1 KY1 HOF
FCoV2
55-67 kDa
1:2000 1:50 1:50 1:50
UGA4.4 KM1 KN1 KY1 KY2 HOF
SCoV2
32-37 kDa
1:50 1:50 1:50

### S6 Fig. SCoV2 Wuhan *versus* SCoV1 Tor2 Spike Glycoproteins

## A

```
                                Signal Peptide → NTD / S1
YP_009724390.1_SCoV2_Wuhan      MFVFLVLLPLVSSQCVNLTT--RTQLPPAY--TNSFTRGVYYPDKVFRSSVLHSTQDLFL
YP_009825051.1_SCoV1_Tor2       MFIFLLFLTLTSGSDLDRCTTFDDVQAPNYTQHTSSMRGVYYPDEIFRSDTLYLTQDLFL
                                **:**::* *.*.. ::  *        * *    .*  *******::***..*: ******

YP_009724390.1_SCoV2_Wuhan      PFFSNVTWFHAIHVSGTNGTKRFDNPVLPFNDGVYFASTEKSNIIRGWIFGTTLDSKTQS
YP_009825051.1_SCoV1_Tor2       PFYSNVTGFHTIN-------HTFGNPVIPFKDGIYFAATEKSNVVRGWVFGSTMNNKSQS
                                **:**** **:*.        : * ***:**:**:***:*****::***:**:*::.*:**

YP_009724390.1_SCoV2_Wuhan      LLIVNNATNVVIKVCEFQFCNDPFLGVYYHKNNKSWMESEFRVYSSANNCTFEYVSQPFL
YP_009825051.1_SCoV1_Tor2       VIIINNSTNVVIRACNFELCDNPFFAVSKPMGT----QTHTMIFDNAFNCTFEYISDAFS
                                ::*:**:*****:.*:*::*::**:.*      .     ::.  ::..* *******:*: *

YP_009724390.1_SCoV2_Wuhan      MDLEGKQGNFKNLREFVFKNIDGYFKIYSKHTPINLVRDLPQGFSALEPLVDLPIGINIT
YP_009825051.1_SCoV1_Tor2       LDVSEKSGNFKHLREFVFKNKDGFLYVYKGYQPIDVVRDLPSGFNTLKPIFKLPLGINIT
                                :*:. *.****.******** **:: :*. : **::*****.**.:*:*:..**:*****

YP_009724390.1_SCoV2_Wuhan      RFQTLLALHRSYLTPGDSSSGWTAGAAAYYVGYLQPRTFLLKYNENGTITDAVDCALDPL
YP_009825051.1_SCoV1_Tor2       NFRAILTAFS------PAQDIWGTSAAAYFVGYLKPTTFMLKYDENGTITDAVDCSQNPL
                                .*:::*: .         :.. * :.****:****:* **:***:***********: :**

                                NTD ←                  → RBD / S1
YP_009724390.1_SCoV2_Wuhan      SETKCTLKSFTVEKGIYQTSNFRVQPTESIVRFPNITNLCPFGEVFNATRFASVYAWNRK
YP_009825051.1_SCoV1_Tor2       AELKCSVKSFEIDKGIYQTSNFRVVPSGDVVRFPNITNLCPFGEVFNATKFPSVYAWERK
                                :* **::*** ::********** *: .:*******************:* *****:**

YP_009724390.1_SCoV2_Wuhan      RISNCVADYSVLYNSASFSTFKCYGVSPTKLNDLCFTNVYADSFVIRGDEVRQIAPGQTG
YP_009825051.1_SCoV1_Tor2       KISNCVADYSVLYNSTFFSTFKCYGVSATKLNDLCFSNVYADSFVVKGDDVRQIAPGQTG
                                :**************: ********** ********:********::**:**********

YP_009724390.1_SCoV2_Wuhan      KIADYNYKLPDDFTGCVIAWNSNNLDSKVGGNYNYLYRLFRKSNLKPFERDISTEIYQAG
YP_009825051.1_SCoV1_Tor2       VIADYNYKLPDDFMGCVLAWNTRNIDATSTGNYNYKYRYLRHGKLRPFERDISNVPFSPD
                                 ************ ***:***:.*:*:.   ***** **  :*:.:*:*******.  :.

                                                                               RBD ←
YP_009724390.1_SCoV2_Wuhan      STPCNGVEGFNCYFPLQSYGFQPTNGVGYQPYRVVVLSFELLHAPATVCGPKKSTNLVKN
YP_009825051.1_SCoV1_Tor2       GKPCTP-PALNCYWPLNDYGFYTTTGIGYQPYRVVVLSFELLNAPATVCGPKLSTDLIKN
                                ..**.   .:***:**:.***  *.*:***************.********* **:*:**

                                   → CTD / S1
YP_009724390.1_SCoV2_Wuhan      KCVNFNFNGLTGTGVLTESNKKFLPFQQFGRDIADTTDAVRDPQTLEILDITPCSFGGVS
YP_009825051.1_SCoV1_Tor2       QCVNFNFNGLTGTGVLTPSSKRFQPFQQFGRDVSDFTDSVRDPKTSEILDISPCAFGGVS
                                :**************** *.*:* ********::* **:****:* *****:**:*****

YP_009724390.1_SCoV2_Wuhan      VITPGTNTSNQVAVLYQDVNCTEVPVAIHADQLTPTWRVYSTGSNVFQTRAGCLIGAEHV
YP_009825051.1_SCoV1_Tor2       VITPGTNASSEVAVLYQDVNCTDVSTAIHADQLTPAWRIYSTGNNVFQTQAGCLIGAEHV
                                *******:*.:***********:*  .*********:**:****.*****:**********

                                                     CTD ←
                                                  ← S1/S2 →
                                                         ↓
YP_009724390.1_SCoV2_Wuhan      NNSYECDIPIGAGICASYQTQTNSPRRARSVASQSIIAYTMSLGAENSVAYSNNSIAIPT
YP_009825051.1_SCoV1_Tor2       DTSYECDIPIGAGICASYHTVSL----LRSTSQKSIVAYTMSLGADSSIAYSNNTIAIPT
                                :.****************:* :        **.:.:**:********:.*:*****:*****

YP_009724390.1_SCoV2_Wuhan      NFTISVTTEILPVSMTKTSVDCTMYICGDSTECSNLLLQYGSFCTQLNRALTGIAVEQDK
YP_009825051.1_SCoV1_Tor2       NFSISITTEVMPVSMAKTSVDCNMYICGDSTECANLLLQYGSFCTQLNRALSGIAAEQDR
                                **:**:***::****:******.**********:*****************:***.***:

                                                               S2'
                                                                ↓
YP_009724390.1_SCoV2_Wuhan      NTQEVFAQVKQIYKTPPIKDFGGFNFSQILPDPSKPSKRSFIEDLLFNKVTLADAGFIKQ
YP_009825051.1_SCoV1_Tor2       NTREVFAQVKQMYKTPTLKYFGGFNFSQILPDPLKPTKRSFIEDLLFNKVTLADAGFMKQ
                                **:********:****  :* ************* **:********************:**
```

```
YP_009724390.1_SCoV2_Wuhan      YGDCLGDIAARDLICAQKFNGLTVLPPLLTDEMIAQYTSALLAGTITSGWTFGAGAALQI
YP_009825051.1_SCoV1_Tor2       YGECLGDINARDLICAQKFNGLTVLPPLLTDDMIAAYTAALVSGTATAGWTFGAGAALQI
                                **:***** **********************:*** **:**::** *:************

YP_009724390.1_SCoV2_Wuhan      PFAMQMAYRFNGIGVTQNVLYENQKLIANQFNSAIGKIQDSLSSTASALGKLQDVVNQNA
YP_009825051.1_SCoV1_Tor2       PFAMQMAYRFNGIGVTQNVLYENQKQIANQFNKAISQIQESLTTTSTALGKLQDVVNQNA
                                ************************* ******.**.:**:**::*::*************

YP_009724390.1_SCoV2_Wuhan      QALNTLVKQLSSNFGAISSVLNDILSRLDKVEAEVQIDRLITGRLQSLQTYVTQQLIRAA
YP_009825051.1_SCoV1_Tor2       QALNTLVKQLSSNFGAISSVLNDILSRLDKVEAEVQIDRLITGRLQSLQTYVTQQLIRAA
                                ************************************************************

YP_009724390.1_SCoV2_Wuhan      EIRASANLAATKMSECVLGQSKRVDFCGKGYHLMSFPQSAPHGVVFLHVTYVPAQEKNFT
YP_009825051.1_SCoV1_Tor2       EIRASANLAATKMSECVLGQSKRVDFCGKGYHLMSFPQAAPHGVVFLHVTYVPSQERNFT
                                **************************************:**************:**:***

YP_009724390.1_SCoV2_Wuhan      TAPAICHDGKAHFPREGVFVSNGTHWFVTQRNFYEPQIITTDNTFVSGNCDVVIGIVNNT
YP_009825051.1_SCoV1_Tor2       TAPAICHEGKAYFPREGVFVFNGTSWFITQRNFFSPQIITTDNTFVSGNCDVVIGIINNT
                                *******:***:******** *** **:*****:.*********************:***

YP_009724390.1_SCoV2_Wuhan      VYDPLQPELDSFKEELDKYFKNHTSPDVDLGDISGINASVVNIQKEIDRLNEVAKNLNES
YP_009825051.1_SCoV1_Tor2       VYDPLQPELDSFKEELDKYFKNHTSPDVDLGDISGINASVVNIQKEIDRLNEVAKNLNES
                                ************************************************************

YP_009724390.1_SCoV2_Wuhan      LIDLQELGKYEQYIKWPWYIWLGFIAGLIAIVMVTIMLCCMTSCCSCLKGCCSCGSCCKF
YP_009825051.1_SCoV1_Tor2       LIDLQELGKYEQYIKWPWYVWLGFIAGLIAIVMVTILLCCMTSCCSCLKGACSCGSCCKF
                                *******************:****************:*************.*********

YP_009724390.1_SCoV2_Wuhan      DEDDSEPVLKGVKLHYT
YP_009825051.1_SCoV1_Tor2       DEDDSEPVLKGVKLHYT
                                *****************
```

## B

##### Summary of Amino Acid (AA) Sequence Identity and Similarity between SCoV2 and SCoV1

| | **NTD** | **RBD** | **CTD** | **S1** | **S2** | **Spike** |
|---|---|---|---|---|---|---|
| Total No. of AA | 322 | 211 | 144 | 676 | 588 | 1277 |
| No. of AA with Identity | 164 | 154 | 111 | 438 | 529 | 967 |
| No. of AA with Similarity | 259 | 190 | 132 | 594 | 579 | 1173 |
| % Identity | 50.9% | 73.0% | 77.1% | 64.8% | 90.0% | 75.7% |
| % Similarity | **80.4%** | 90.0% | 91.7% | 87.9% | **98.5%** | 91.9% |

#### S7 Fig. SCoV2 UF2-RBD Sequence with AA Extensions

**A** **RTFLLKYNENGTITDAVDCALDPLSETKCTLKSFTVEKGIYQTSNF**RVQPTESIVRFPNITNLCPFGEVFNAT
RFASVYAWNRKRISNCVADYSVLYNSASFSTFKCYGVSPTKLNDLCFTNVYADSFVIRGDEVRQIAPGQTGKI
ADYNYKLPDDFTGCVIAWNSNNLDSKVGGNYNYLYRLFRKSNLKPFERDISTEIYQAGSTPCNGVEGFNCYFP
LQSYGFQPTNGVGYQPYRVVVLSFELLHAPATVCGPKK**STNLVKNKCVNF**

**B** **SCoV2 RBD Immunoblot Analysis**

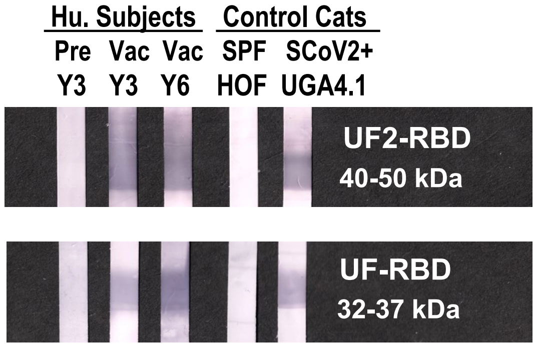


**C** **Description of Human Subjects* and Laboratory Cats**

| Subject or Cat | Age (yr) Gender | Vac Freq | Vaccination Types | SCoV2 Infection |
|---|---|---|---|---|
| Y3 Pre | 63F | - | - | - |
| Y3 Vac | 68F | 3 | Pfizer S mRNA | - |
| Y6 Vac | 38F | 3 | Astra-Zeneca-S<br>Pfizer S mRNA | - |
| UGA4.1 | 1F | - | - | + |
| SPF HOF | 1F | - | - | - |

* Blood from the subjects collected using UF IRB202002902.
** Abbreviations: year (yr), Vaccinated/Vaccination (Vac), frequency (Freq), Pre-vaccination (Pre), female (F), human (Hu), specific pathogen free (SPF)

### S8 Fig. 10 CCoV2 Spike Sequences *versus* Four FCoV2 Spike Sequences

## A

```
                                           Signal Peptide
KP981644_CCoV2a_Italy_CB/05           MIVILTCVLLLCSYHTVASTTNNDCRQVNVTQLDGNENLIRDFLFQ---NFKEEGTVVVG
KC175339_CCoV2_Germany_171            MIVLVTCLLLLCSYHTVLSTTNNECIQVNVTQLAGNENLIRDFLFS---NFKEEGSVVVG
KC175340_CCoV2_US,NY_K378             MIVLILCLL-LFSYNSVICTSNNDCVQVNVTQLPGNENIIKDFLFH---TFKEEGSVVVG
KC175341_CCoV2_US,NY_S378             MIVLILCLL-LFSYNSVICTSNNDCVQVNVTQLPGNENIIKDFLFH---TFKEEGSVVVG
JQ404410_CCoV2_US,GA_UGA-TN449        MIVLVTCILLLCSYHTVLSTANNDCRQVNVTQLPGNEYLIRDFLFQ---SFKEEGSVVVG
JQ404409_CCoV2_US,GA_UGA-1-71         MIVLILCLL-LFSYNSVICTSNNDCVQVNVTQLPGNENIIKDFLFH---TFKEEGSVVVG
KY063616_CCoV2_China_HLJ071           MIVLVTCILLLCSYHTVLSTANNDCRQVNVTQLPGNEYLIRDFLFQ---SFKEEGSVVVG
KY063617_CCoV2_China_HLJ072           MIVLVTCILLLCSYHTVLSTANNDCRQVNVTQLPGNEYLIRDFLFQ---SFKEEGSVVVG
KY063618_CCoV2_China_HLJ073           MIVLVTCILLLCSYHTVLSTANNDCRQVNVTQLPGNEYLIRDFLFQ---SFKEEGSVVVG
GQ477367_CCoV2_Taiwan_NTU336          MKFLVFVVIV-----PLIYGDEFPCSKFINRTIGNHWNLIETFLLNYSSRLPPNSDVVLG
GQ152141_FCoV2_Taiwan_NTU156          -MIVLTLFILLVSYTNVYCTANSDCVQVNVTQLPGNENIIKDFLFS---NFKEEGTVVVG
JQ408981_FCoV2_Hungary_DF2            MIVLVTCLLLLCSYHTVLSTTNNECIQVNVTQLAGNENLIRDFLFS---NFKEEGSVVVG
JN634064_FCoV2_WSU79-1683             MIVLVTCILLLCSYHTVSSTSNNDCRQVNVTQLAGNENLIRDFLFQ---SFKEEGIVVVG
DQ010921_FCoV2_WSU79-1146             MIVLVTCLLLLCSYHTVLSTTNNECIQVNVTQLAGNENLIRDFLFS---NFKEEGSVVVG
                                       .::  .:      :    :  * :.    :  .  :*. **:     :  :. **:*
                                                         NTD / S1

KP981644_CCoV2a_Italy_CB/05           GYYPT-EVWYNCSRTAPTTAYEYFNNIHAFYFDMEAMENSTGNARGKPLLFHVHGEPVSA
KC175339_CCoV2_Germany_171            GYYPT-EVWYNCSRTARTTAFQYFNNIHAFYFVMEAMENSTGNARGKPLLFHVHGEPVSV
KC175340_CCoV2_US,NY_K378             GYYPT-EVWYNCSRSATTTAYKDFSNIHAFYFDMEAMENSTGNARGKPLLVHVHGDPVSI
KC175341_CCoV2_US,NY_S378             GYYPT-EVWYNCSRSATTTAYKDFSNIHAFYFDMEAMENSTGNARGKPLLVHVHGDPVSI
JQ404410_CCoV2_US,GA_UGA-TN449        GYYPT-EVWYNCSRTATTTAYQYFNNIHAFYFDMEAMENSTGNARGKPLLFHVHGDPVSV
JQ404409_CCoV2_US,GA_UGA-1-71         GYYPT-EVWYNCSRSATTTAYKDFSNIHAFYFDMEAMENSTGNARGKPLLVHVHGDPVSI
KY063616_CCoV2_China_HLJ071           GYYPT-EVWYNCSRTATTTAYQYFNNIHAFYFDMEAMENSTGNARGKPLLFHVHGDPVSV
KY063617_CCoV2_China_HLJ072           GYYPT-EVWYNCSRTATTTAYQYFNNIHAFYFDMEAMENSTGNARGKPLLFHVHGDPVSV
KY063618_CCoV2_China_HLJ073           GYYPT-EVWYNCSRTATTTAYQYFNNIHAFYFDMEAMENSTGNARGKPLLFHVHGDPVSV
GQ477367_CCoV2_Taiwan_NTU336          DYFPTVTPWFNCIRNNNNSLYVTMENLKALYWDYATENITA-D-HRQRLHVVVQGNPYSI
GQ152141_FCoV2_Taiwan_NTU156          GYYPT-EVWYNCSRTATTTAYEYFSNIHAFYFDMEAMENSTGNARGKPLLFHVHGEPVSV
JQ408981_FCoV2_Hungary_DF2            GYYPT-EVWYNCSRTAQTTAFQYFNNIHAFYFVMEAMENSTGNARGKPLLFHVHGEPVSV
JN634064_FCoV2_WSU79-1683             GYYPT-EVWYNCSRTATTTAYEYFNNIHAFYFDMEAMENSTGNARGKPLLFHVHGEPV--
DQ010921_FCoV2_WSU79-1146             GYYPT-EVWYNCSRTARTTAFQYFNNIHAFYFVMEAMENSTGNARGKPLLFHVHGEPVSV
                                        *:**   *:** *.  .: :  :.*::*:*:   : : :: : : : * . *:*:*

KP981644_CCoV2a_Italy_CB/05           IIYISAYRNDVQHRPLLKHGLVCITKTRN-----IDYNSFTSRQWNSICTGNDR---KIP
KC175339_CCoV2_Germany_171            I--ISAYRDDVQQRPLLKHGLVCITKNRH-----INYEQFTSNQWNSTCTGADR---KIP
KC175340_CCoV2_US,NY_K378             IIYISAYRDDVQGRPLLKHGLLCITKNKI-----IDYNTFTSAQWSAICLGDDR---KIP
KC175341_CCoV2_US,NY_S378             IIYISAYRDDVQGRPLLKHGLLCITKNKI-----IDYNTFTSAQWSAICLGDDR---KIP
JQ404410_CCoV2_US,GA_UGA-TN449        IIYISAYRDDVQHRPLLKHGLVCITKSRN-----IDYNTFTSSQWNSICTGNDR---KIP
JQ404409_CCoV2_US,GA_UGA-1-71         IIYISAYRDDVQGRPLLKHGLLCITKNKI-----IDYNTFTSAQWSAICLGDDR---KIP
KY063616_CCoV2_China_HLJ071           IIYISAYRDDVQHRPLLKHGLVCITKSRN-----IDYNTFTSSQWNSICTGNDR---KIP
KY063617_CCoV2_China_HLJ072           IIYISAYRDDVQHRPLLKHGLVCITKSRN-----IDYNTFTSSQWNSICTGNDR---KIP
KY063618_CCoV2_China_HLJ073           IIYISAYRDDVQHRPLLKHGLVCITKSRN-----IDYNTFTSSQWNSICTGNDR---KIP
GQ477367_CCoV2_Taiwan_NTU336          TVTTTRDFN------SAEGAIICICKGTPPKTTTVNSASQLNCNWGTECRLQHTGTQHTK
GQ152141_FCoV2_Taiwan_NTU156          IIYISAYRDDVQQRPLLKHGLVCITESRN-----IDYNQFTSNQWNSICTGNDR---KIP
JQ408981_FCoV2_Hungary_DF2            IIYISAYRDDVQQRPLLKHGLVCITKNRH-----INYEQFTSNQWNSTCTGADR---KIP
JN634064_FCoV2_WSU79-1683             IIYISAYGDDVQQRPLLEHGLLCITKNRN-----IDYNTFTSNQWDSICTGNDR---KIP
DQ010921_FCoV2_WSU79-1146             I--ISAYRDDVQQRPLLKHGLVCITKNRH-----INYEQFTSNQWNSTCTGADR---KIP
                                          :   :         : .::** :          ::      . :* : *    .    :

KP981644_CCoV2a_Italy_CB/05           F----SVIPTDNGTKIYGLEWNDEFVTAYISGHSYNWNINNNWFNNVTLLYSRSSTATWQ
KC175339_CCoV2_Germany_171            F----SVIPTDNGTKIYGLEWNDDFVTAYISGRSYHLNINTNWFNNVTLLYSRSSTATWE
KC175340_CCoV2_US,NY_K378             F----SVIPTDNGTKIFGLEWNDDYVTAYISDRSHHLNINNNWFNNVTILYSRSSTATWQ
KC175341_CCoV2_US,NY_S378             F----SVIPTDNGTKIFGLEWNDDYVTAYISDRSHHLNINNNWFNNVTILYSRSSTATWQ
JQ404410_CCoV2_US,GA_UGA-TN449        F----SVIPTDNGTKIYGLEWNDEFVTAYISGHSYNWNINNNWFNNVTLLYSRSSTATWQ
JQ404409_CCoV2_US,GA_UGA-1-71         F----SVIPTDNGTKIFGLEWNDDYVTAYISDRSHHLNINNNWFNNVTILYSRSSTATWQ
KY063616_CCoV2_China_HLJ071           F----SVIPTDNGTKIYGLEWNDEFVTAYISGHSYNWNINNNWFNNVTLLYSRSSTATWQ
KY063617_CCoV2_China_HLJ072           F----SVIPTDNGTKIYGLEWNDEFVTAYISGHSYNWNINNNWFNNVTLLYSRSSTATWQ
KY063618_CCoV2_China_HLJ073           F----SVIPTDNGTKIYGLEWNDEFVTAYISGHSYNWNINNNWFNNVTLLYSRSSTATWQ
GQ477367_CCoV2_Taiwan_NTU336          FPICPSNIGSNCGNMLYGLQWFTDEVVAYLHGAVYRINFENQWSGTVTLGDMRATTAQTA
GQ152141_FCoV2_Taiwan_NTU156          F----SVIPTDNGTKIYGLEWNDESVTAYISGHSYSLNINTNWFNNVTLLYSRSSTATWQ
JQ408981_FCoV2_Hungary_DF2            F----SVIPTDNGTKIYGLEWNDDFVTAYISGRSYHLNINTNWFNNVTLLYSRSSTATWE
JN634064_FCoV2_WSU79-1683             F----SVIPRDNGTKIYGLEWNDEFVTAYISGRSYNWNINNNWFNNVTLLYSRSSTATWH
DQ010921_FCoV2_WSU79-1146             F----SVIPTDNGTKIYGLEWNDDFVTAYISGRSYHLNINTNWFNNVTLLYSRSSTATWE
                                      *    * *  : *. ::**:*  : *.**:    :  *::.:*  .**:   *::**
```

```
KP981644_CCoV2a_Italy_CB/05        --HSAAYVYQGVSNFTYYKLNNTNGLKTYEFCDDYEYCTGYATNVFAPTVGGYIPDGFSF
KC175339_CCoV2_Germany_171         --YSAAYAYQGVSNFTYYKLNNTNGLKTYELCEDYEHCTGYATNVFAPTSGGYIPDGFSF
KC175340_CCoV2_US,NY_K378          --KSAAYVYQGVSNFTYYKLNNTNGLKSYELCEDYEYCTGYATNVFAPTVGGYIPDGFSF
KC175341_CCoV2_US,NY_S378          --KSAAYVYQGVSNFTYYKLNNTNGLKSYELCEDYEYCTGYATNVFAPTVGGYIPDGFSF
JQ404410_CCoV2_US,GA_UGA-TN449     --HSAAYVYQGVSNFTYYKLNNTNGLKTYEFCEDYEYCTATATNVFAPTVGGYIPDGFSF
JQ404409_CCoV2_US,GA_UGA-1-71      --KSAAYVYQGVSNFTYYKLNNTNGLKSYELCEDYEYCTGYATNVFAPTVGGYIPDGFSF
KY063616_CCoV2_China_HLJ071        --HSAAYVYQGVSNFTYYKLNNTNGLKTYEFCEDYEYCTGHATNVFAPTVGGYIPDGFSF
KY063617_CCoV2_China_HLJ072        --HSAAYVYQGVSNFTYYKLNNTNGLKTYEFCEDYEYCTGHATNVFAPTVGGYIPDGFSF
KY063618_CCoV2_China_HLJ073        --HSAAYVYQGVSNFTYYKLNNTNGLKTYEFCEDYEYCTGHATNVFAPTVGGYIPDGFSF
GQ477367_CCoV2_Taiwan_NTU336       GAVVDLWWFNPVYDITYYRVNSKNSTTVIFNC--TDQCASYVDNIFTTQPGGFIPSDFSF
GQ152141_FCoV2_Taiwan_NTU156       --KSAAYVYQGVSNFTYYKLNNTNGLKTYELCENYEYCTGHATNVFAPIAGGYIPDGFSF
JQ408981_FCoV2_Hungary_DF2         --YSAAYAYQGVSNFTYYKLNNTNGLKTYELCEDYEHCTGYATNVFAPTSGGYIPDGFSF
JN634064_FCoV2_WSU79-1683          --HSAAYVYQGVSNFTYYKLNNTNGLKTYEFCEDYEYCT-YATNVFAPTVGGYIPDGFSF
DQ010921_FCoV2_WSU79-1146          --YSAAYAYQGVSNFTYYKLNNTNGLKTYELCEDYEHCTGYATNVFAPTSGGYIPDGFSF
                                         : :: * ::***::*..*. .     *   : *:  . *:*:   **:**. ***

KP981644_CCoV2a_Italy_CB/05        NNWFLLTNGSTFVSGRFVTNQPLLVNCLWPVPSFGVAAQEFCFEGAQFSQCNGVSLNNTV
KC175339_CCoV2_Germany_171         NNWFLLTNSSTFVSGRFVTNQPLLINCLWPVPSFGVAAQEFCFEGAQFSQCNGVSLNNTV
KC175340_CCoV2_US,NY_K378          NNWFMLTNSSTFVSGRFVTNQPLLVNCLWPVPSFGVAAQQFCFEGAQFSQCNGVSLNNTV
KC175341_CCoV2_US,NY_S378          NNWFMLTNSSTFVSGRFVTNQPLLVNCLWPVPSFGVAAQQFCFEGAQFSQCNGVSLNNTV
JQ404410_CCoV2_US,GA_UGA-TN449     NNWFLLTNSSTFVSGRFVTNQPLLVNCLWPVPSFGVAAQEFCFEGAQFSQCNGMSLNNTV
JQ404409_CCoV2_US,GA_UGA-1-71      NNWFMLTNSSTFVSGRFVTNQPLLVNCLWPVPSFGVAAQQFCFEGAQFSQCNGVSLNNTV
KY063616_CCoV2_China_HLJ071        NNWFLLTNSSTFVSGRFVTNQPLLVNCLWPVPSFGVAAQEFCFEGAQFSQCNGMSLNNTV
KY063617_CCoV2_China_HLJ072        NNWFLLTNSSTFVSGRFVTNQPLLVNCLWPVPSFGVAAQEFCFEGAQFSQCNGMSLNNTV
KY063618_CCoV2_China_HLJ073        NNWFLLTNSSTFVSGRFVTNQPLLVNCLWPVPSFGVAAQEFCFEGAQFSQCNGMSLNNTV
GQ477367_CCoV2_Taiwan_NTU336       NNWFLLTNSSTVLNGKLVTRQPLLVNCLWPVPSFKGAASTFCFEGAGFDQCNGAVLNNTV
GQ152141_FCoV2_Taiwan_NTU156       NNWFLLTNSSTFVSGRFVTNQPLLVNCLWPVPSFGVAAQEFCFEGAQFSQCNGVSLNNTV
JQ408981_FCoV2_Hungary_DF2         NNWFLLTNSSTFVSGRFVTNQPLLINCLWPVPSFGVAAQEFCFEGAQFSQCNGVSLNNTV
JN634064_FCoV2_WSU79-1683          NNWFLLTNSSTFVSGRFVTNQPLLVNCLWPVPSFGVAAQEFCFEGAQFSQCSGVSLNNTV
DQ010921_FCoV2_WSU79-1146          NNWFLLTNSSTFVSGRFVTNQPLLINCLWPVPSFGVAAQEFCFEGAQFSQCNGVSLNNTV
                                   ****:***.**.:.*::**.****:*********  **. ****** *.**.*  *****

KP981644_CCoV2a_Italy_CB/05        DVIRFNLNFTADVQSGMGATVFSLNTTGGVILEISCYNDTVSESSFYSYGEIPFGITDGP
KC175339_CCoV2_Germany_171         DVIRFNLNFTADVQSGMGATVFSLNTTGGVILEISCYSDTVSESSSYSYGEIPFGITDGP
KC175340_CCoV2_US,NY_K378          DVIRFNLNFTTDVQSGMGATVFSLNTTGGVILEISCYNDTVSESSFYSYGEISFGVTDGP
KC175341_CCoV2_US,NY_S378          DVIRFNLNFTTDVQSGMGATVFSLNTTGGVILEISCYNDTVSESSFYSYGEISFGVTDGP
JQ404410_CCoV2_US,GA_UGA-TN449     DVIRFNLNFTADVQSGMGATVFSLNTTGGVILEISCYKDTVSESSFYSYGEIPFGITDGP
JQ404409_CCoV2_US,GA_UGA-1-71      DVIRFNLNFTTDVQSGMGATVFSLNTTGGVILEISCYNDTVSESSFYSYGEISFGVTDGP
KY063616_CCoV2_China_HLJ071        DVIRFNLNFTADVQSGMGATVFSLNTTGGVILEISCYKDTVSESSFYSYGEIPFGITDGP
KY063617_CCoV2_China_HLJ072        DVIRFNLNFTADVQSGMGATVFSLNTTGGVILEISCYKDTVSESSFYSYGEIPFGITDGP
KY063618_CCoV2_China_HLJ073        DVIRFNLNFTADVQSGMGATVFSLNTTGGVILEISCYKDTVSESSFYSYGEIPFGITDGP
GQ477367_CCoV2_Taiwan_NTU336       DVIRFNLNFTADVQSGMGATVFSLNTTGGVILEISCYNDTVSESSFYSYGEIPFGVTDGP
GQ152141_FCoV2_Taiwan_NTU156       DVIRFNLNLTADVQSGMGATVFSLNTTGGVILEISCYNDTVSESSFYSYGEIPFGVTDGP
JQ408981_FCoV2_Hungary_DF2         DVIRFNLNFTADVQSGMGATVFSLNTTGGVILEISCYSDTVSESSSYSYGEIPFGITDGP
JN634064_FCoV2_WSU79-1683          DVIRSILISLQ-MYNLVWVLQYFRRYTGGVILEISCYNDTVSESSFYSYGEIPFGITDGP
DQ010921_FCoV2_WSU79-1146          DVIRFNLNFTADVQSGMGATVFSLNTTGGVILEISCYSDTVSESSSYSYGEIPFGITDGP
                                   ****  *      : . : .  :  . ***********.******* ****** **:****
                                                                                      NTD

KP981644_CCoV2a_Italy_CB/05        RYCYALYNGTALKYLGTLPPSVKEIAISKWGHFYINGYNFFSTFPIDCISFNLTTGTSGA
KC175339_CCoV2_Germany_171         RYCYVLYNGTALKYLGTLPPSVKEIAISKWGHFYINGYNFFSTFPIGCISFNLTTGVSGA
KC175340_CCoV2_US,NY_K378          RYCFALYNGTALKYLGTLPPSVKEIAISKWGHFYINGYNFFSTFPIDCISFNLTTGDSGA
KC175341_CCoV2_US,NY_S378          RYCFALYNGTALKYLGTLPPSVKEIAISKWGHFYINGYNFFSTFPIDCISFNLTTGDSGA
JQ404410_CCoV2_US,GA_UGA-TN449     RYCYVLYNGTALKYLGTLPPSVKEIAISKWGHFYINGYNFFSTFPIDCISFNLTTGASGA
JQ404409_CCoV2_US,GA_UGA-1-71      RYCYALYNGTALKYLGTLPPSVKEIAISKWGHFYINGYNFFSTFPIDCISFNLTTGDSGA
KY063616_CCoV2_China_HLJ071        RYCYVLYNGTALKYLGTLPPSVKEIAISKWGHFYINGYNFFSTFPIDCISFNLTTGASGA
KY063617_CCoV2_China_HLJ072        RYCYVLYNGTALKYLGTLPPSVKEIAISKWGHFYINGYNFFSTFPIDCISFNLTTGASGA
KY063618_CCoV2_China_HLJ073        RYCYVLYNGTALKYLGTLPPSVKEIAISKWGHFYINGYNFFSTFPIDCISFNLTTGASGA
GQ477367_CCoV2_Taiwan_NTU336       RYCYVLYNGTALKYLGTLPPSVKEIAISKWGHFYINGYNFFSTFPIDCISFNLTTGASGA
GQ152141_FCoV2_Taiwan_NTU156       RYCYVLYNGTALKYLGTLPPSVKEIAISKWGHFYINGYNFFSTFPIDCISFNLTTGASGA
JQ408981_FCoV2_Hungary_DF2         RYCYVLYNGTALKYLGTLPPSVKEIAISKWGHFYINGYNFFSTFPIGCISFNLTTGASGA RBD
JN634064_FCoV2_WSU79-1683          RYCYVLYNGTALKYLGTLPPSVKEIAISKWGHFYINGYNFFSTFPIDCISFNLTTGDSGA 53T
DQ010921_FCoV2_WSU79-1146          RYCYVLYNGTALKYLGTLPPSVKEIAISKWGHFYINGYNFFSTFPIGCISFNLTTGVSGA 51I
                                   ***:.****************************************** ********* *** 51S
                                          Proposed RBD / S1
```

```
KP981644_CCoV2a_Italy_CB/05        FWTIAYTSYTEALVQVENTAIKKVTYCNSHINNIKCSQLTANLQNGFYPVASSEVGLVNK
KC175339_CCoV2_Germany_171         FWTIAYTSYTEALVQVENTAIKNVTYCNSHINNIKCSQLTANLNNGFYPVASSEVGFVNK
KC175340_CCoV2_US,NY_K378          FWTIAYTSYTDALVQVENTAIKKVTYCNSHINNIKCSQLTANLQNGFYPVASSEVGLVNK
KC175341_CCoV2_US,NY_S378          FWTIAYTSYTDALVQVENTAIKKVTYCNSHINNIKCSQLTANLQNGFYPVASSEVGLVNK
JQ404410_CCoV2_US,GA_UGA-TN449     FWTIAYTSYTEALVQVENTAIKKVTYCNSHINNIKCSQLTANLNNGFYPVASSEVGLVNK
JQ404409_CCoV2_US,GA_UGA-1-71      FWTIAYTSYTDALVQVENTAIKKVTYCNSHINNIKCSQLTANLQNGFYPVASSEVGLVNK
KY063616_CCoV2_China_HLJ071        FWTIAYTSYTEALVQVENTAIKKVTYCNSHINNIKCSQLTANLNNGFYPVASSEVGLVNK
KY063617_CCoV2_China_HLJ072        FWTIAYTSYTEALVQVENTAIKKVTYCNSHINNIKCSQLTANLNNGFYPVASSEVGLVNK
KY063618_CCoV2_China_HLJ073        FWTIAYTSYTEALVQVENTAIKKVTYCNSHINNIKCSQLTANLNNGFYPVASSEVGLVNK
GQ477367_CCoV2_Taiwan_NTU336       FWTIAYTSYTEALVQVENTAIKKVTYCNSHINNIKCSQLTANLQNGFYPVASSEVGLVNK
GQ152141_FCoV2_Taiwan_NTU156       FWTIAYTSYTEALVQVENTAIKKVTYCNSHINNIKCSQLTANLQNGFYPVASSDVGLVNK
JQ408981_FCoV2_Hungary_DF2         FWTIAYTSYTEALVQVENTAIKNVTYCNSHINNIKCSQLTANLNNGFYPVASSEVGFVNK RBD
JN634064_FCoV2_WSU79-1683          FWTIAYTSYTEALVQVENTAIKKVTYCNSHINNIKCSQLTANLNNGFYPVASSEVGLVNK 60T
DQ010921_FCoV2_WSU79-1146          FWTIAYTSYTEALVQVENTAIKNVTYCNSHINNIKCSQLTANLNNGFYPVASSEVGFVNK 55I
                                   **********:***********:*******************:*********:**:*** 60S

KP981644_CCoV2a_Italy_CB/05        SVVLLPSFYSHTSVNITIDLGMKRSGYGQPIASALSNITLPMQDNNTDVYCIRSNQFSVY
KC175339_CCoV2_Germany_171         SVVLLPSFFTYTAVNITIDLGMKLSGYGQPIASTLSNITLPMQDNNTDVYCIRSNQFSVY
KC175340_CCoV2_US,NY_K378          SVVLLPSFYSHTSVNITIDLGMKRSGYGQPIASTLSNITLPMQDNNTDVYCIRSNRFSVY
KC175341_CCoV2_US,NY_S378          SVVLLPSFYSHTSVNITIDLGMKRSGYGQPIASTLSNITLPMQDNNTDVYCIRSNRFSVY
JQ404410_CCoV2_US,GA_UGA-TN449     SVVLLPTFFAHTAFNITIDLGMKRSGYGQPIASTLSNITLPMQDNNTDVYCVRSNQFSVY
JQ404409_CCoV2_US,GA_UGA-1-71      SVVLLPSFYSHTSVNITIDLGMKRSGYGQPIASTLSNITLPMQDNNTDVYCIRSNQFSVY
KY063616_CCoV2_China_HLJ071        SVVLLPTFFAHTAFNITIDLGMKRSGYGQPIASTLSNITLPMQDNNTDVYCVRSNQFSVY
KY063617_CCoV2_China_HLJ072        SVVLLPTFFAHTAFNITIDLGMKRSGYGQPIASTLSNITLPMQDNNTDVYCVRSNQFSVY
KY063618_CCoV2_China_HLJ073        SVVLLPTFFAHTAFNITIDLGMKRSGYGQPIASTLSNITLPMQDNNTDVYCVRSNQFSVY
GQ477367_CCoV2_Taiwan_NTU336       SVVLLPSFFAHTAVNITIDLGMKRSGYGQPIASTLSNITLPMQDNNTDVYCIRSNQFSVY
GQ152141_FCoV2_Taiwan_NTU156       SVVLLPIFFAHTAVNITIDLGMKRSGYGQPIASTLSNITLPMQDNNTDVYCVRSNQFSVY
JQ408981_FCoV2_Hungary_DF2         SVVLLPSFFTHTAVNITIDLGMKLSGYGQPIASTLSNITLPMQDNNTDVYCIRSNQFSVY RBD
JN634064_FCoV2_WSU79-1683          SVVLLPIFFAHTAINITIDLGMKRSGYGQPIASTLSNITLPMQDNNTDVYCIRSNQFSVY 60T
DQ010921_FCoV2_WSU79-1146          SVVLLPSFFTYTAVNITIDLGMKLSGYGQPIASTLSNITLPMQDNNTDVYCIRSNQFSVY 50I
                                   ****** *:::*:.********* *********:****************:***:**** 58S

KP981644_CCoV2a_Italy_CB/05        VHSTCKSSLWDNVFNSDCTDFLHATAVIKTGTCPFSFDKLNNYLTFNKFCFSLNPVGANC
KC175339_CCoV2_Germany_171         VHSTCKSSLWDNIFNQDCTDVLEATAVIKTGTCPFSFDKLNNYLTFNKFCLSLSPVGANC
KC175340_CCoV2_US,NY_K378          FHSTCKSSLWDDVFNSDCTDVLYATAVIKTGTCPFSFDKLNNYLTFNKFCLSLNPVGANC
KC175341_CCoV2_US,NY_S378          FHSTCKSSLWDDVFNSDCTDVLYATAVIKTGTCPFSFDKLNNYLTFNKFCLSLNPVGANC
JQ404410_CCoV2_US,GA_UGA-TN449     VHSTCKSSSWDNIFNQDCTDVLEATAVIKTGTCPFSFDKLNNYLTFNKFCLSLSPVGANC
JQ404409_CCoV2_US,GA_UGA-1-71      VHSTCKSSLWDDVFNSDCTDVLYATAVIKTGTCPFSFDKLNNYLTFNKFCLSLNPVGANC
KY063616_CCoV2_China_HLJ071        VHSTCKSSSWDNIFNQDCTDVLEATAVIKTGTCPFSFDKLNNYLTFNKFCLSLSPVGANC
KY063617_CCoV2_China_HLJ072        VHSTCKSSSWDNIFNQDCTDVLEATAVIKTGTCPFSFDKLNNYLTFNKFCLSLSPVGANC
KY063618_CCoV2_China_HLJ073        VHSTCKSSSWDNIFNQDCTDVLEATAVIKTGTCPFSFDKLNNYLTFNKFCLSLSPVGANC
GQ477367_CCoV2_Taiwan_NTU336       THSTCKSSLWDNNFNRDCTDVLYATAVIKTGTCPFSFDKLNNYLTFNKLCLSLNPTGANC
GQ152141_FCoV2_Taiwan_NTU156       THSTCKSSLWDNNFNQDCTDVLYATAVIKTGTCPFSFDKLNNYLTFNKLCLSLNPTGANC
JQ408981_FCoV2_Hungary_DF2         VPSTCKSSLWDNIFNQDCTDVLEATAVIKTGTCPFSFDKLNNYLTFNKFCLSLSPVGANC RBD
JN634064_FCoV2_WSU79-1683          VHSICKSSLWDNIFNQECTDVLDATAVIKTGTCPFSFDKLNNYLTFNKFCLSLSPVGANC 60T
DQ010921_FCoV2_WSU79-1146          VHSTCKSSLWDNIFNQDCTDVLEATAVIKTGTCPFSFDKLNNYLTFNKFCLSLSPVGANC 46I
                                     * **** **: ** :***.* *************************:*:**.*.**** 53S

KP981644_CCoV2a_Italy_CB/05        KFDVAARTRTNEQVVRSLYVIYEEGDNIVGVPSDNSGLHDLSVLHLDSCTDYNIYGRTGV
KC175339_CCoV2_Germany_171         KFDVAARTRTNEQVVRSLYVIYEEGDNIVGVPSDNSGLHDLSVLHLDSCTDYNIYGRTGV
KC175340_CCoV2_US,NY_K378          KFDVAARTRTNEQVVRSLYVIYEEGDNIVGVPSDNSGLHDLSVLHLDSCTDYNIYGITGV
KC175341_CCoV2_US,NY_S378          KFDVAARTRTNEQVVRSLYVIYEEGDNIVGVPSDNSGLHDLSVLHLDSCTDYNIYGITGV
JQ404410_CCoV2_US,GA_UGA-TN449     KFDVAAHTRTNEQVVRSLYVIYEEGDNIVGVPSDNSGLHDLSVLHLDSCTDYNIYGRTGV
JQ404409_CCoV2_US,GA_UGA-1-71      KFDVAARTRTNEQVVRSLYVIYEEGDNIVGVPSDNSGLHDLSVLHLDSCTDYNIYGRTGV
KY063616_CCoV2_China_HLJ071        KFDVAAHTRTNEQVVRSLYVIYEEGDNIVGVPSDNSGLHDLSVLHLDSCTDYNIYGRTGV
KY063617_CCoV2_China_HLJ072        KFDVAAHTRTNEQVVRSLYVIYEEGDNIVGVPSDNSGLHDLSVLHLDSCTDYNIYGRTGV
KY063618_CCoV2_China_HLJ073        KFDVAAHTRTNEQVVRSLYVIYEEGDNIVGVPSDNSGLHDLSVLHLDSCTDYNIYGRTGV
GQ477367_CCoV2_Taiwan_NTU336       KFDVAARTRTSEQVVRSLYVIYEEGDNIVGVPSDNSGLHDLSVLHLDSCTDYNIYGRTGV
GQ152141_FCoV2_Taiwan_NTU156       KFDVAARTRTNEQVVRSLYVIYEEGDNIVGVPSDNSGPHDLSVLHLDSCTDYNIYGRTGV
JQ408981_FCoV2_Hungary_DF2         KFDVAARTRTNEQVVRSLYVIYEEGDNIVGVPSDNSGLHDLSVLHLDSCTDYNIYGRTGV RBD
JN634064_FCoV2_WSU79-1683          KFDVAARTRTNEQVVRSLYVIYEEGDNIVGVPSDNSGLHDLSVLHLDSCTEYNIYGRTGV 35T
DQ010921_FCoV2_WSU79-1146          KFDVAARTRTNEQVVRSLYVIYEEGDNIVGVPSDNSGLHDLSVLHLDSCTDYNIYGRTGV 33I
                                   ******:***.************************** ************:******** 35S
```

Proposed RBD

CTD / S1

RBD Total (T): 268 aa
RBD Identity (I): 235 / 268 = 87.7%
RBD Similarity (S): 257 / 268 = 95.9 %

```
KP981644_CCoV2a_Italy_CB/05       GIIRQTNSTLLSGLYYTSLSGDLLGFKNVTDGVVYSVTPCDVSAQAAVIDGAIVGAMTSI
KC175339_CCoV2_Germany_171        GIIRRTNSTLLSGLYYTSLSGDLLGFKNVSDGVIYSVTPCDVSAQAAVIDGAIVGAMTSI
KC175340_CCoV2_US,NY_K378         GIIRQTNSTLLSGLYYTSLSGDLLGFKNVSDGVIYSVTPCDVSAHAAVIDGAIVGAMTSI
KC175341_CCoV2_US,NY_S378         GIIRQTNSTLLSGLYYTSLSGDLLGFKNVSDGVIYSVTPCDVSAHAAVIDGAIVGAMTSI
JQ404410_CCoV2_US,GA_UGA-TN449    GIIRQTNSTLLSGLYYTSLSGDLLGFKNVSDGVIYSVTPCDVSAQVAVIDGAIVGAMTSI
JQ404409_CCoV2_US,GA_UGA-1-71     GIIRQTNSTLLSGLYYTSLSGDLLGFKNVSDGVIYSVTPCDVSAQAAVIDGAIVGAMTSI
KY063616_CCoV2_China_HLJ071       GIIRQTNSTLLSGLYYTSLSGDLLGFKNVSDGVIYSVTPCDVSAQVAVIDGAIVGAMTSI
KY063617_CCoV2_China_HLJ072       GIIRQTNSTLLSGLYYTSLSGDLLGFKNVSDGVIYSVTPCDVSAQVAVIDGAIVGAMTSI
KY063618_CCoV2_China_HLJ073       GIIRQTNSTLLSGLYYTSLSGDLLGFKNVSDGVIYSVTPCDVSAQVAVIDGAIVGAMTSI
GQ477367_CCoV2_Taiwan_NTU336      GIIRQTNSTILSGLYYTSLSGDLLGFKNVSDGVVYSVTPCDVSAQAAVIDGAIVGAMTSI
GQ152141_FCoV2_Taiwan_NTU156      GIIRQTNSTILSGLYYTSLSGDLLGFKNVSDGVVYSVTPCDVSAQAAVIDGAIVGAMTSI
JQ408981_FCoV2_Hungary_DF2        GIIRRTNSTLLSGLYYTSLSGDLLGFKNVSDGVIYSVTPCDVSAQAAVIDGAIVGAMTSI
JN634064_FCoV2_WSU79-1683         GIIRQTNSTLLSGLYYTSLSGDLLGFKNVSDGVIYSVTPCDVSAQAAVIDGAIVGAMTSI
DQ010921_FCoV2_WSU79-1146         GIIRRTNSTLLSGLYYTSLSGDLLGFKNVSDGVIYSVTPCDVSVQAAVIDGAIVGAMTSI
                                  ****:****:*******************:***:*********.:.**************

KP981644_CCoV2a_Italy_CB/05       NSELLGLTHWTTTPNFYYYSIYNYTNARTRGTAIDSNDVDCEPIITYSNIGVCKNGALVF
KC175339_CCoV2_Germany_171        NSELLGLTHWTTTPNFYYYSIYNYTSERTRGTAIDSNDVDCEPVITYSNIGVCKNGALVF
KC175340_CCoV2_US,NY_K378         NSELLGLTHWTTTPNFYYYSIYNYTNERTRGTAIDSNDVDCEPIITYSNIGVCKNGALVF
KC175341_CCoV2_US,NY_S378         NSELLGLTHWTTTPNFYYYSIYNYTNERTRGTAIDSNDVDCEPIITYSNIGVCKNGALVF
JQ404410_CCoV2_US,GA_UGA-TN449    NSELLGLTHWTTTPNFYYYSIYNYTNERTRGTAIDSNDVDCEPIITYANIGVCKNGALVF
JQ404409_CCoV2_US,GA_UGA-1-71     NSELLGLTHWTTTPNFYYYSIYNYTNERTRGTAIDSNDVDCEPIITYSNIGVCKNGALVF
KY063616_CCoV2_China_HLJ071       NSELLGLTHWTTTPNFYYYSIYNYTNERTRGTAIDSNDVDCEPIITYANIGVCKNGALVF
KY063617_CCoV2_China_HLJ072       NSELLGLTHWTTTPNFYYYSIYNYTNERTRGTAIDSNDVDCEPIITYANIGVCKNGALVF
KY063618_CCoV2_China_HLJ073       NSELLGLTHWTTTPNFYYYSIYNYTNERTRGTAIDSNDVDCEPIITYANIGVCKNGALVF
GQ477367_CCoV2_Taiwan_NTU336      NSELLGLTHWTTTPNFYYYSIYNYTNERTRGTAIDSNDVDCEPIITYSNIGVCKNGALVF
GQ152141_FCoV2_Taiwan_NTU156      NSELLGLTHWTTTPNFYYYSIYNYTNERTRGTAIDSNAVDCEPIITYSNIGVCKNGALVF
JQ408981_FCoV2_Hungary_DF2        NSELLGLTHWTTTPNFYYYSIYNYTSERTRGTAIDSNDVDCEPVITYSNIGVCKNGALVF
JN634064_FCoV2_WSU79-1683         NSELLGLKHWTTTPNFYYYSIYNYTNERTRGTAIDSNDVDCEPIITYSNIGVCKNGALVF
DQ010921_FCoV2_WSU79-1146         NSELLGLTHWTTTPNFYYYSIYNYTSERTRGTAIDSNDVDCEPVITYSNIGVCKNGALVF
                                  *******.*****************. ********** *****:***:************

KP981644_CCoV2a_Italy_CB/05       INVTHSDGDVQPISTGNVTIPTNFTISVQVEYIQVYTTPVSIDCSRYVCNGNPRCNKLLT
KC175339_CCoV2_Germany_171        INVTHSDGDVQPISTGNVTIPTNFTISVQVEYMQVYTTPVSIDCARYVCNGNPRCNKLLT
KC175340_CCoV2_US,NY_K378         INVTHSDGDVQPISTGNVTIPTNFTISVQVEYIQVYTTPVSIDCSRYVCNGNPRCNKLLT
KC175341_CCoV2_US,NY_S378         INVTHSDGDVQPISTGNVTIPTNFTISVQVEYIQVYTTPVSIDCSRYVCNGNPRCNKLLT
JQ404410_CCoV2_US,GA_UGA-TN449    INVTHSDGDVQPISTGNVTIPTNFTISVQVEYIQVYTTPVSIDCSRYVCNGNPRCNKLLT
JQ404409_CCoV2_US,GA_UGA-1-71     INVTHSDGDVQPISTGNVTIPTNFTISVQVEYIQVYTTPVSIDCSRYVCNGNPRCNKLLT
KY063616_CCoV2_China_HLJ071       INVTHSDGDVQPISTGNVTIPTNFTISVQVEYIQVYTTPVSIDCSRYVCNGNPRCNKLLT
KY063617_CCoV2_China_HLJ072       INVTHSDGDVQPISTGNVTIPTNFTISVQVEYIQVYTTPVSIDCSRYVCNGNPRCNKLLT
KY063618_CCoV2_China_HLJ073       INVTHSDGDVQPISTGNVTIPTNFTISVQVEYIQVYTTPVSIDCSRYVCNGNPRCNKLLT
GQ477367_CCoV2_Taiwan_NTU336      INVTHSDGDVQPISTGNVTIPTNFTISVQVEYIQVYTTPVSIDCSRYVCNGNPRCNKLLT
GQ152141_FCoV2_Taiwan_NTU156      INVTHSDGDVQPISTGNVTIPTNFTISVQVEYIQVYTTPVSIDCSRYVCNGNPRCNKLLT
JQ408981_FCoV2_Hungary_DF2        INVTHSDGDVQPISTGNVTIPTNFTISVQVEYMQVYTTPVSIDCARYVCNGNPRCNKLLT
JN634064_FCoV2_WSU79-1683         INVTHSDGDVQPISTGTVTIPTNFTISVQVEYIQVYTTPVSIDCARYVCNGNPRCNKLLT
DQ010921_FCoV2_WSU79-1146         INVTHSDGDVQPISTGNVTIPTNFTISVQVEYMQVYTTPVSIDCARYVCNGNPRCNKLLT
                                  ****************.***************:***********:***************

KP981644_CCoV2a_Italy_CB/05       QYVSACQTIEQALAMGARLENMEVDSMLFVSENALKLASVEAFNSTETLDPIYKEWPNIG
KC175339_CCoV2_Germany_171        QYVSACQTIEQALAMGARLENMEVDSMLFVSENALKLASVEAFNSTENLDPIYKEWPSIG
KC175340_CCoV2_US,NY_K378         QYVSACQTIEQALAMGARLENMEIDSMLFVSENALKLASVEAFNSTETLDPIYKEWPNIG
KC175341_CCoV2_US,NY_S378         QYVSACQTIEQALAMGARLENMEIDSMLFVSENALKLASVEAFNSTETLDPIYKEWPNIG
JQ404410_CCoV2_US,GA_UGA-TN449    QYVSACQTIEQALAMGARLENMEVDSMLFVSENALKLASVEAFNSTENLDPIYKEWPNIG
JQ404409_CCoV2_US,GA_UGA-1-71     QYVSACQTIEQALAMGARLENMEIDSMLFVSENALKLASVEAFNSTETLDPIYKEWPNIG
KY063616_CCoV2_China_HLJ071       QYVSACQTIEQALAMGARLENMEVDSMLFVSENALKLASVEAFNSTENLDPIYKEWPNIG
KY063617_CCoV2_China_HLJ072       QYVSACQTIEQALAMGARLENMEVDSMLFVSENALKLASVEAFNSTENLDPIYKEWPNIG
KY063618_CCoV2_China_HLJ073       QYVSACQTIEQALAMGARLENMEVDSMLFVSENALKLASVEAFNSTENLDPIYKEWPNIG
GQ477367_CCoV2_Taiwan_NTU336      QYVSACQTIEQALAMSARLENMEVDSMLFVSENALKLASVEAFNSTEHLDPIYKEWPNIG
GQ152141_FCoV2_Taiwan_NTU156      QYVSACQTIEQALAMSARLENMEVDSMLFVSENALKLASVEAFNSTEHLDPIYKEWPNIG
JQ408981_FCoV2_Hungary_DF2        QYVSACQTIEQALAMGARLENMEVDSMLFVSENALKLASVEAFNSTENLDPIYKEWPSIG
JN634064_FCoV2_WSU79-1683         QYVSACQTIEQALAMGARLENMEVDSMLFVSENALKLASVEAFNSTENLDPIYKEWPNIG
DQ010921_FCoV2_WSU79-1146         QYVSACQTIEQALAMGARLENMEVDSMLFVSENALKLASVEAFNSTENLDPIYKEWPSIG
                                  ***************.*******:*********************** *********.**
```

```
KP981644_CCoV2a_Italy_CB/05        GSWLGGLKDILPSHNSKRKYRSAIEDLLFDKVVTSGLGTVDEDYKRCTGGYDIADLVCAQ
KC175339_CCoV2_Germany_171         GSWLGGLKDILPSHNSKRKYGSAIEDLLFDKVVTSGLGTVDEDYKRCTGGYDIADLVCAQ
KC175340_CCoV2_US,NY_K378          GSWLGGLKDILPSHNSKRKYRSAIEDLLFDKVVTSGLGTVDEDYKRCTGGYDIADLVCAQ
KC175341_CCoV2_US,NY_S378          GSWLGGLKDILPSHNSKRKYRSAIEDLLFDKVVTSGLGTVDEDYKRCTGGYDIADLVCAQ
JQ404410_CCoV2_US,GA_UGA-TN449     GSWLGGLKDILPSHNSKRKYRSAIEDLLFDKVVTSGLGTVDEDYKRCTGGYDIADLVCAQ
JQ404409_CCoV2_US,GA_UGA-1-71      GSWLGGLKDILPSHNSKRKYRSAIEDLLFDKVVTSGLGTVDEDYKRCTGGYDIADLVCAQ
KY063616_CCoV2_China_HLJ071        GSWLGGLKDILPSHNSKRKYRSAIEDLLFDKVVTSGLGTVDEDYKRCTGGYDIADLVCAQ
KY063617_CCoV2_China_HLJ072        GSWLGGLKDILPSHNSKRKYRSAIEDLLFDKVVTSGLGTVDEDYKRCTGGYDIADLVCAQ
KY063618_CCoV2_China_HLJ073        GSWLGGLKDILPSHNSKRKYRSAIEDLLFDKVVTSGLGTVDEDYKRCTGGYDIADLVCAQ
GQ477367_CCoV2_Taiwan_NTU336       GSWLGGLKDILPSHNSKRKYRSAIEDLLFDKVVTSGLGTVDEDYKRCTGGYDIADLVCAQ
GQ152141_FCoV2_Taiwan_NTU156       GSWLGGLKDILPSHNSKRKYRSAIEDLLFDKVVTSGLGTVDEDYKRCTGGYDIADLVCAQ
JQ408981_FCoV2_Hungary_DF2         GSWLGGLKDILPSHNSKRKYGSAIEDLLFDKVVTSGLGTVDEDYKRCTGGYDIADLVCAQ
JN634064_FCoV2_WSU79-1683          GSWLGGLKDILPSHNSKRKYRSAIEDLLFDKVVTSGLGTVDEDYKRCTGGYDIADLVCAQ
DQ010921_FCoV2_WSU79-1146          GSWLGGLKDILPSHNSKRKYGSAIEDLLFDKVVTSGLGTVDEDYKRCTGGYDIADLVCAQ
                                   ******************** ***************************************
```

S1/S2

CTD

```
KP981644_CCoV2a_Italy_CB/05        YYNGIMVLPGVANDDKMAMYTASLAGGITLGALGGGAVSIPFAVAVQARLNYVALQTDVL
KC175339_CCoV2_Germany_171         YYNGIMVLPGVANADKMTMYTASLAGGITLGALGGGAVAIPFAVAVQARLNYVALQTDVL
KC175340_CCoV2_US,NY_K378          YYNGIMVLPGVANDDKMAMYTASLAGGITLGSLGGGAVSIPFAIAVQARLNYVALQTDVL
KC175341_CCoV2_US,NY_S378          YYNGIMVLPGVANDDKMAMYTASLAGGITLGSLGGGAVSIPFAIAVQARLNYVALQTDVL
JQ404410_CCoV2_US,GA_UGA-TN449     YYNGIMVLPGVANDDKMTMYTASLAGGITLGALGGGAVAIPFAVAVQARLNYVALQTDVL
JQ404409_CCoV2_US,GA_UGA-1-71      YYNGIMVLPGVANDDKMAMYTASLAGGITLGALGGGAVSIPFAIAVQARLNYVALQTDVL
KY063616_CCoV2_China_HLJ071        YYNGIMVLPGVANDDKMTMYTASLAGGITLGALGGGAVAIPFAVAVQARLNYVALQTDVL
KY063617_CCoV2_China_HLJ072        YYNGIMVLPGVANDDKMTMYTASLAGGITLGALGGGAVAIPFAVAVQARLNYVALQTDVL
KY063618_CCoV2_China_HLJ073        YYNGIMVLPGVANDDKMTMYTASLAGGITLGALGGGAVAIPFAVAVQARLNYVALQTDVL
GQ477367_CCoV2_Taiwan_NTU336       YYNGIMVLPGVANDDKMAMYTASLAGGITLGALGGGAVSIPFAVAVQARLNYVALQTDVL
GQ152141_FCoV2_Taiwan_NTU156       YYNGIMVLPGVANDDKMTMYTASLAGGITLGALGGGAVAIPFAVAVQARLNYVALQTDVL
JQ408981_FCoV2_Hungary_DF2         YYNGIMVLPGVANADKMTMYTASLAGGITLGALGGGAVAIPFAVAVQARLNYVALQTDVL
JN634064_FCoV2_WSU79-1683          YYNGIMVLPGVANDDKMTMYTASLAGGITLGALGGGAVAIPFAVAVQARLNYVALQTDVL
DQ010921_FCoV2_WSU79-1146          YYNGIMVLPGVANADKMTMYTASLAGGITLGALGGGAVAIPFAVAVQARLNYVALQTDVL
                                   ************* ***:*************:*******:***:****************

KP981644_CCoV2a_Italy_CB/05        NKNQQILANAFNQAIGNITQAFGKVNDAIHQTSKGLATVAKALAKVQDVVNTQGQALSHL
KC175339_CCoV2_Germany_171         NKNQQILANAFNQAIGNITQAFGKVNDAIHQTSQGLATVAKALAKVQDVVNTQGQALSHL
KC175340_CCoV2_US,NY_K378          NKNQQILANAFNQAIGNITQAFGKVNDAIHQTSQGLATVAKVLAKVQDVVNTQGQALSHL
KC175341_CCoV2_US,NY_S378          NKNQQILANAFNQAIGNITQAFGKVNDAIHQTSQGLATVAKVLAKVQDVVNTQGQALSHL
JQ404410_CCoV2_US,GA_UGA-TN449     NKNQQILANAFNQAIGNITQAFGKVNDAIHQTSQGLATVAKALAKVQDVVNTQGQALSHL
JQ404409_CCoV2_US,GA_UGA-1-71      NKNQQILANAFNQAIGNITQAFGKVNDAIHQTSQGLATVAKALAKVQDVVNTQGQALSHL
KY063616_CCoV2_China_HLJ071        NKNQQILANAFNQAIGNITQAFGKVNDAIHQTSQGLATVAKALAKVQDVVNTQGQALSHL
KY063617_CCoV2_China_HLJ072        NKNQQILANAFNQAIGNITQAFGKVNDAIHQTSQGLATVAKALAKVQDVVNTQGQALSHL
KY063618_CCoV2_China_HLJ073        NKNQQILANAFNQAIGNITQAFGKVNDAIHQTSQGLATVAKALAKVQDVVNTQGQALSHL
GQ477367_CCoV2_Taiwan_NTU336       NKNQQILANAFNQAIGNITQAFGKVNDAIHQTSQGLATVAKALAKVQDVVNTQGQALSHL
GQ152141_FCoV2_Taiwan_NTU156       NKNQQILANAFNQAIGNITQAFGKVNDAIHQTSQGLATVAKALVKVQDVVNTQGQALSHL
JQ408981_FCoV2_Hungary_DF2         NKNQQILANAFNQAIGNITQAFGKVNDAIHQTSQGLATVAKALAKVQDVVNTQGQALSHL
JN634064_FCoV2_WSU79-1683          NKNQQILANAFNQAIGNITQAFGKVNDAIHQTSKGLATVAKALAKVQDVVNTQGQALSHL
DQ010921_FCoV2_WSU79-1146          NKNQQILANAFNQAIGNITQAFGKVNDAIHQTSQGLATVAKALAKVQDVVNTQGQALSHL
                                   *********************************:*******.*.****************

KP981644_CCoV2a_Italy_CB/05        TVQLQNNFQAISSSISDIYNRLDELSADAQVDRLITGRLTALNAFVSQTLTRQAEVRASR
KC175339_CCoV2_Germany_171         TVQLQNNFQAISSSISDIYNRLDELSADAQVDRLITGRLTALNAFVSQTLTRQAEVRASR
KC175340_CCoV2_US,NY_K378          TLQLQNNFQAISSSISDIYNRLDELSADAQVDRLITGRLTALNAFVSQTLTRQAEVRASR
KC175341_CCoV2_US,NY_S378          TVQLQNNFQAISSSISDIYNRLDELSADAQVDRLITGRLTALNAFVSQTLTRQAEVRASR
JQ404410_CCoV2_US,GA_UGA-TN449     TVQLQNNFQAISSSISDIYNRLDELSADAQVDRLITGRLTALNAFVSQTLTRQAEVRASR
JQ404409_CCoV2_US,GA_UGA-1-71      TVQLQNNFQAISSSISDIYNRLDELSADAQVDRLITGRLTALNAFVSQTLTRQAEVRASR
KY063616_CCoV2_China_HLJ071        TVQLQNNFQAISSSISDIYNRLDELSADAQVDRLITGRLTALNAFVSQTLTRQAEVRASR
KY063617_CCoV2_China_HLJ072        TVQLQNNFQAISSSISDIYNRLDELSADAQVDRLITGRLTALNAFVSQTLTRQAEVRASR
KY063618_CCoV2_China_HLJ073        TVQLQNNFQAISSSISDIYNRLDELSADAQVDRLITGRLTALNAFVSQTLTRQAEVRASR
GQ477367_CCoV2_Taiwan_NTU336       TVQLQNNFQAISSSISDIYNRLDELSADAQVDRLITGRLTALNAFVSQTLTRQAEVRASR
GQ152141_FCoV2_Taiwan_NTU156       TVQLQNNFQAISSSISDIYNRLDELSADAQVDRLITGRLTALNAFVSQTLTRQAEVRASR
JQ408981_FCoV2_Hungary_DF2         TVQLQNNFQAISSSISDIYNRLDELSADAQVDRLITGRLTALNAFVSQTLTRQAEVRASR
JN634064_FCoV2_WSU79-1683          TVQLQNNFQAISSSISDIYNRLDELSADAQVDRLITGRLTALNAFVSQTLTRQAEVRASR
DQ010921_FCoV2_WSU79-1146          TVQLQNNFQAISSSISDIYNRLDELSADAQVDRLITGRLTALNAFVSQTLTRQAEVRASR
                                   *:**********************************************************
```

```
KP981644_CCoV2a_Italy_CB/05       QLAKDKVNECVRSQSQRFGFCGNGTHLFSLANAAPNGMIFFHTVLLPTAYETVTAWSGIC
KC175339_CCoV2_Germany_171        QLAKDKVNECVRSQSQRFGFCGNGTHLFSLANAAPNGMIFFHTVLLPTAYETVTAWSGIC
KC175340_CCoV2_US,NY_K378         QLAKDKVNECVRSQSQRFGFCGNGTHLFSLANAAPNGMIFFHTVLLPTAYETVTAWSGIC
KC175341_CCoV2_US,NY_S378         QLAKDKVNECVRSQSQRFGFCGNGTHLFSLANAAPNGMIFFHTVLLPTAYETVTAWSGIC
JQ404410_CCoV2_US,GA_UGA-TN449    QLAKDKVNECVRSQSQRFGFCGNGTHLFSLANAAPNGMIFFHTVLLPTAYETVTAWSGIC
JQ404409_CCoV2_US,GA_UGA-1-71     QLAKDKVNECVRSQSQRFGFCGNGTHLFSLANAAPNGMIFFHTVLLPTAYETVTAWSGIC
KY063616_CCoV2_China_HLJ071       QLAKDKVNECVRSQSQRFGFCGNGTHLFSLANAAPNGMIFFHTVLLPTAYETVTAWSGIC
KY063617_CCoV2_China_HLJ072       QLAKDKVNECVRSQSQRFGFCGNGTHLFSLANAAPNGMIFFHTVLLPTAYETVTAWSGIC
KY063618_CCoV2_China_HLJ073       QLAKDKVNECVRSQSQRFGFCGNGTHLFSLANAAPNGMIFFHTVLLPTAYETVTAWSGIC
GQ477367_CCoV2_Taiwan_NTU336      QLAKDKVNECVRSQSQRFGFCGNGTHLFSLANAAPNGMIFFHSVLLPTAYETVTAWSGIC
GQ152141_FCoV2_Taiwan_NTU156      QLAKDKVNECVRSQSQRFGFCGNGTHLFSLANAAPNGMIFFHTVLLPTAYETVTAWSGIC
JQ408981_FCoV2_Hungary_DF2        QLAKDKVNECVRSQSQRFGFCGNGTHLFSLANAAPNGMIFFHTVLLPTAYETVTAWSGIC
JN634064_FCoV2_WSU79-1683         QLAKDKVNECVRSQSQRFGFCGNGTHLFSLANAAPNGMIFFHTVLLPTAYETVTAWPGIC
DQ010921_FCoV2_WSU79-1146         QLAKDKVNECVRSQSQRFGFCGNGTHLFSLANAAPNGMIFFHTVLLPTAYETVTAWSGIC
                                  ******************************************:************* ***

KP981644_CCoV2a_Italy_CB/05       ASDGDRTFGLVVKDVQLTLFRNLDDKFYLTPRTMYQPRVATSSDFVQIEGCDVLFVNATV
KC175339_CCoV2_Germany_171        ASDGDRTFGLVVKDVQLTLFRNLDDKFYLTPRTMYQPRVATSSDFVQIEGCDVLFVNATV
KC175340_CCoV2_US,NY_K378         ASDGDRTFGLVVKDVQLTLFRNLDDKFYLTPRTMYQPRVATSSDFVQIEGCDVLFVNATV
KC175341_CCoV2_US,NY_S378         ASDGDRTFGLVVKDVQLTLFRNLDDKFYLTPRTMYQPRVATSSDFVQIEGCDVLFVNATV
JQ404410_CCoV2_US,GA_UGA-TN449    ASDGDRTFGLVVKDVQLTLFRNLDGKFYLTPRTMYQPRVVTSSDFVQIEGCDVLFVNATV
JQ404409_CCoV2_US,GA_UGA-1-71     ASDGDRTFGLVVKDVQLTLFRNLDDKFYLTPRTMYQPRVATSSDFVQIEGCDVLFVNATV
KY063616_CCoV2_China_HLJ071       ASDGDRTFGLVVKDVQLTLFRNLDGKFYLTPRTMYQPRVATSSDFVQIEGCDVLFVNATV
KY063617_CCoV2_China_HLJ072       ASDGDRTFGLVVKDVQLTLFRNLDGKFYLTPRTMYQPRVATSSDFVQIEGCDVLFVNATV
KY063618_CCoV2_China_HLJ073       ASDGDRTFGLVVKDVQLTLFRNLDGKFYLTPRTMYQPRVATSSDFVQIEGCDVLFVNATV
GQ477367_CCoV2_Taiwan_NTU336      ASDGDRTFGLVVKDVQLTLFRNLDGKFYLTPRTMYQPRAATSSDFVQIEGCDVLFVNATV
GQ152141_FCoV2_Taiwan_NTU156      ASDGDRTFGLVVKDVQLTLFRNLDDKFYLTPRTMYQPRAATSSDFVQIEGCDVLFVNATV
JQ408981_FCoV2_Hungary_DF2        ASDGDRTFGLVVKDVQLTLFRNLDDKFYLTPRTMYQPRVATSSDFVQIEGCDVLFVNATV
JN634064_FCoV2_WSU79-1683         ASDGDRTFGLVVKDVQLTLFRNLDDKFYLTPRTMYQPRAATSSDFVQIEGCDVLFVNATV
DQ010921_FCoV2_WSU79-1146         ASDGDRTFGLVVKDVQLTLFRNLDDKFYLTPRTMYQPRVATSSDFVQIEGCDVLFVNATV
                                  ************************ *************..********************

KP981644_CCoV2a_Italy_CB/05       IELPSIIPDYIDINQTVQDILENYRPNWTVPELTLDIFNATYLNLTGEINDLEFRSEKLH
KC175339_CCoV2_Germany_171        IDLPSIIPDYIDINQTVQDILENYRPNWTVPEFTLDIFNATYLNLTGEIDDLEFRSEKLH
KC175340_CCoV2_US,NY_K378         IDLPSIIPDYIDINQTVQDILENFRPNWTVPELPLDIFNATYLNLTGEINDLEFRSEKLH
KC175341_CCoV2_US,NY_S378         IDLPSIIPDYIDINQTVQDILENFRPNWTVPELPLDIFNATYLNLTGEINDLEFRSEKLH
JQ404410_CCoV2_US,GA_UGA-TN449    IDLPSIIPDYIDINQTVQDILENYRPNWTVPELTLDIFNATYLNLTGEIDDLEFRSEKLH
JQ404409_CCoV2_US,GA_UGA-1-71     IDLPSIIPDYIDINQTVQDILENFRPNWTVPELPLDIFNATYLNLTGEINDLEFRSEKLH
KY063616_CCoV2_China_HLJ071       IDLPSIIPDYIDINQTVQDILENYRPNWTVPELTLDIFNATYLNLTGEIDDLEFRSEKLH
KY063617_CCoV2_China_HLJ072       IDLPSIIPDYIDINQTVQDILENYRPNWTVPELTLDIFNATYLNLTGEIDDLEFRSEKLH
KY063618_CCoV2_China_HLJ073       IDLPSIIPDYIDINQTVQDILENYRPNWTVPELTLDIFNATYLNLTGEIDDLEFRSEKLH
GQ477367_CCoV2_Taiwan_NTU336      IDLPSIIPDYIDINQTVQDILENYRPNWTVPELTLDIFNATYLNLTGEIDDLEFRSEKLH
GQ152141_FCoV2_Taiwan_NTU156      IDLPSIIPDYIDINQTVQDILENYRPNWTVPELTLDIFNATYLNLTGEIDDLEFRSEKLH
JQ408981_FCoV2_Hungary_DF2        IDLPSIIPDYIDINQTVQDILENYRPNWTVPEFTLDIFNATYLNLTGEIDDLEFRSEKLH
JN634064_FCoV2_WSU79-1683         IDLPSIIPDYIDINQTVQDILENYRPNWTVPELTLDIFNATYLNLTGEIDDLEFRSEKLH
DQ010921_FCoV2_WSU79-1146         IDLPSIIPDYIDINQTVQDILENYEPNWTVPEFTLDIFNATYLNLTGEIDDLEFRSEKLH
                                  *:*********************:.*******: ***************:**********

KP981644_CCoV2a_Italy_CB/05       NTTVELAVLIDNINNTLVNLEWLNRIETYVKWPWYVWLLIGLVVIFCIPLLLFCCCSTGC
KC175339_CCoV2_Germany_171        NTTVELAILIDNINNTLVNLEWLNRIETYVKWPWYVWLLIGLVVVFCIPLLLFCCFSTGC
KC175340_CCoV2_US,NY_K378         NTTVELAILIDNINNTLVNLEWLNRIETYVKWPWYVWLLIGLVVIFCIPILLFCCCSTGC
KC175341_CCoV2_US,NY_S378         NTTVELAILIDNINNTLVNLEWLNRIETYVKWPWYVWLLIGLVVIFCIPILLFCCCSTGC
JQ404410_CCoV2_US,GA_UGA-TN449    NTTVELAILIDNINNTLVNLEWLNRIETYVKWPWYVWLLIGLVVIFCIPLLLFCCCSTGC
JQ404409_CCoV2_US,GA_UGA-1-71     NTTVELAILIDNINNTLVNLEWLNRIETYVKWPWYVWLLIGLVVIFCIPILLFCCCSTGC
KY063616_CCoV2_China_HLJ071       NTTVELAILIDNINNTLVNLEWLNRIETYVKWPWYVWLLIGLVVIFCIPLLLFCCCSTGC
KY063617_CCoV2_China_HLJ072       NTTVELAILIDNINNTLVNLEWLNRIETYVKWPWYVWLLIGLVVIFCIPLLLFCCCSTGC
KY063618_CCoV2_China_HLJ073       NTTVELAILIDNINNTLVNLEWLNRIETYVKWPWYVWLLIGLVVIFCIPLLLFCCCSTGC
GQ477367_CCoV2_Taiwan_NTU336      NTTVELAILIDNINNTLVNLEWLNRIETYVKWPWYVWLLIGLVVVFCIPLLLFCCCSTGC
GQ152141_FCoV2_Taiwan_NTU156      NTTVELAILIDNINNTLVNLEWLNRIETYVKWPWYVWLLIGLVVVFCIPLLLFCCCSTGC
JQ408981_FCoV2_Hungary_DF2        NTTVELAILIDTINNTLVNLEWLNRIETYVKWPWYVWLLIGLVVVFCIPLLLFCCFSTGC
JN634064_FCoV2_WSU79-1683         NTTVELAILIDNINNTLVNLEWLNRIETYVKWPWYVWLLIGLVVIFCIPLLLFCCCSTGC
DQ010921_FCoV2_WSU79-1146         NTTVELAILIDNINNTLVNLEWLNRIETYVKWPWYVWLLIGLVVVFCIPLLLFCCFSTGC
                                  *******:***.********************************:****:***** ****
```

```
KP981644_CCoV2a_Italy_CB/05         CGCIGCLGSCCHSICSRRQFENYEPIEKVHVH
KC175339_CCoV2_Germany_171          CGCIGCLGSCCHSICSRRQFENYEPIEKVHVH
KC175340_CCoV2_US,NY_K378           CGCIGCLGSCCHSICSRRQFESYEPIEKVHVH
KC175341_CCoV2_US,NY_S378           CGCIGCLGSCCHSICSRRQFESYEPIEKVHVH
JQ404410_CCoV2_US,GA_UGA-TN449      CGCIGCLGSCCHSICSRRQFENYEPIEKVHVH
JQ404409_CCoV2_US,GA_UGA-1-71       CGCIGCLGSCCHSICSRRQFESYEPIEKVHVH
KY063616_CCoV2_China_HLJ071         CGCIGCLGSCCHSICSRRQFENYEPIEKVHVH
KY063617_CCoV2_China_HLJ072         CGCIGCLGSCCHSICSRRQFENYEPIEKVHVH
KY063618_CCoV2_China_HLJ073         CGCIGCLGSCCHSICSRRQFENYEPIEKVHVH
GQ477367_CCoV2_Taiwan_NTU336        CGCIGCLGSCCHSICSRRQFENYEPIEKVHVH
GQ152141_FCoV2_Taiwan_NTU156        CGCIGCLGSCCHSICSSRQFENYEPIEKVHVH
JQ408981_FCoV2_Hungary_DF2          CGCIGCLGSCCHSICSRRQFEYYEPIEKVHVH
JN634064_FCoV2_WSU79-1683           CGCIGCLGSCCHSMCSRRQFENYEPIEKVHVH
DQ010921_FCoV2_WSU79-1146           CGCIGCLGSCCHSICSRRQFENYEPIEKVHVH
                                    *************:** **** **********
```

# B

**Summary of Amino Acid (AA) Sequence Identity and Similarity between CCoV2 and FCoV2**

| | **NTD** | **RBD** | **CTD (KRK)** | **CTD (KRKYR)** | **S1 (KRK)** | **S1 (KRKYR)** | **S2 (KRK)** | **S2 (KRKYR)** |
|---|---|---|---|---|---|---|---|---|
| Total No. of AA | 390 | 268 | 267 | 269 | 960 | 962 | 493 | 491 |
| No. of AA with Identity | 127 | 235 | 246 | 247 | 637 | 638 | 464 | 463 |
| No. of AA with Similarity | 229 | 257 | 264 | 265 | 782 | 783 | 485 | 484 |
| % Identity | 32.6% | 87.7% | 92.1% | 91.8% | 66.3 % | 66.3% | 94.1% | 94.3% |
| % Similarity | **58.7%** | 95.9% | **98.9%** | 98.5% | 81.6% | 81.4% | 98.4 % | **98.6%** |

### S9 Fig. Two CCoV1 Spike Sequences *versus* 10 FCoV1 Spike Sequences

## A

```
                                   Signal Peptide
KP849472.1_CCoV1_Italy-23/03       MKIVLFITLLSVVSCDEESYEAQPNDNPAKFNIS-SNSRFEANMYNFLQTWDIPPGTETS
AY307020.1_CCoV1_Elmo/02           MKIFLLSALLAIANCKDE---AGPFDDPQHFNTSFSNQRFEASMYNFLKSWNIPPDTETI
AAB47503_FCoV1_KU-2                -MIFIILTLLSVAKSEDA----PHGVTLPQFNTSHNNERFELNFYNFLQTWDIPPNTETI
KX722530_FCoV1_Dutch_cat2          -MILLILALLSTAKSEDA----PHGVTLPYFNTSYDNNKFELNFYNFLQTWDIPPNTETI
KX722529_FCoV1_Belgium_UG-FH8      -MILLLLAILSIAKCDDA----PNGAILPHFNTSYNNAKFELNFYNFLQTWNIPPNTETI
KP143512_FCoV1_UK26M               --MILLLVLFSVVGAHDA----PHGVTLPQFNTSYNNDKFELNFYNFLQTWDIPPNTETI
MG893511_FCoV1_Germany_Felix       -MIVLIFAILSTALSEDA----PQGVTLPQFNTSYDNSKFELSFYNFLQTWDIPPNTETI
KY566209_FCoV1_China10             -MILLLLAFFSVVKSQDA----PHGVTLPQFNTSHNNDKFELNLFNFLQTWDIPPGTETI
FJ938054_FCoV1_Utrecht_UU4         -MILLILALLSIASSEDA----PHGVTLPHFNTSHNNSKFELNFYNFLQTWDIPPNTETI
DQ848678_FCoV1_C1Je_Kitten_FIP     --MILLLALFSVVSSRDT----PYRVTLPQFNTSHNNDKFELNLFNFLQTWDIPPSTETI
EU186072_FCoV1_Black               -MIVLIFALLSTARSEDA----PHGVTLPQFNTSSNNQKFELNFYNFLQTWDIPPNTETI
AB088222_FCoV1_UCD1                -MILLIFALFSVVRSEDA----PHGVTLPHFNTSYNNPKFELNFYNFLQTWDIPPNTETI
                                     :.:: .::: . . :             ** * .* :** .::***::*:*** ***
                                                  └─► NTD / S1

KP849472.1_CCoV1_Italy-23/03       FGRVIYHYCGQSVNSNTDCAWYNLRWQPG---SGFKEKFLSSAGLGITNVHGFYFDVREH
AY307020.1_CCoV1_Elmo/02           LGGYL-PYCGS----SQRCGWINLLGSDKPNANNKKIQYFDSVSMGITDVHGVYFDVREH
AAB47503_FCoV1_KU-2                LGGYL-PYCGA----GVNCGWYNFS--QSVG-QNGKYAYINTQNLNIPNVHGVYFDVREH
KX722530_FCoV1_Dutch_cat2          LGGYL-PYCGN----GVNCGWYNFVYGRPVG-SNGKYAYINTQNLNIPNVHGVYFDVREH
KX722529_FCoV1_Belgium_UG-FH8      LGGYL-PYCGR----GDNCGWYNFVYQSQGG-TYAKYKYINTEY-LIPNVHGVYFDVREY
KP143512_FCoV1_UK26M               LGGYL-PYCGT----GANCGWYNFVYQQNVG-SNGKYSYINTQNLNIPNVHGVYFDVREH
MG893511_FCoV1_Germany_Felix       LGGYL-PYCGK----GLNCGWYNFVYQQPVT--DAKYSYINTQNLNIPNVHGVYFDVREH
KY566209_FCoV1_China10             LGGYL-PYCGV----GNNCGWYNFVFSQNVG-SNGKYSYINTQNLNIPNVHGVYFDVREH
FJ938054_FCoV1_Utrecht_UU4         LGGYL-PYCDH----EDNCGWYNFVYNNKVG-PNAKYSYINTQNLNIPNVHGVYFDVREH
DQ848678_FCoV1_C1Je_Kitten_FIP     LGGYL-PYCGL----GDNCGWFNFVYKQHDD-IEAKYSYINTQVLNIPNVHGVYFDVREH
EU186072_FCoV1_Black               LGGYL-PYCGD----GPNCGWYNFVYSQNVG-QNGRHSYINTKNLNIPNVHGVYFDVREH
AB088222_FCoV1_UCD1                LGGYL-PYCRT----GQNCGWYNFVYNHAV--QNAKYSYINTQNLNIPNVHGVYFDVREH
                                   :*   :  **          *.* *:            :  ::.:    * :***.******:

KP849472.1_CCoV1_Italy-23/03       D-ENNRWETWDRVGLAVTIYGSSYYDLFMILEDGFSDDSLSIAVKICHWTSGNST---SL
AY307020.1_CCoV1_Elmo/02           AADNNQWDDQERVGLAIVIYGSSWYDLYMLHGNNVESGSLSVSVKICHWTHGSINDTKWY
AAB47503_FCoV1_KU-2                -NNDGEWDDRDKVGLLIAIHGNSKYSLLMVLQDAVEANQPHVAVKICHWK--PGNI---S
KX722530_FCoV1_Dutch_cat2          -NSDGVWDTRDRVGLLIAIHGSAHYSLLMVLQDSVEENQPHFAVKICHWK--PGNI---S
KX722529_FCoV1_Belgium_UG-FH8      GNSDGAWTDMDKVGLLIAIHGTSHYSLLMVLQDRVEENKPHVAVKICHWK--PGNI---S
KP143512_FCoV1_UK26M               -NSDGVWDPRDRVGLLIAIHGKSQYSLLMVLQDNVEENQPHVAVKICHWK--PGNI---S
MG893511_FCoV1_Germany_Felix       -DADGVWETRDRVGLLIAIHGTSHYSLLMVLQDGVEENQPHVAVKICHWK--PGNI---S
KY566209_FCoV1_China10             -NSDGLWDERDRVGLLMSVHGKSHYSLLMVLQDAVEANAPDVAVKICNWQHLTGNI---S
FJ938054_FCoV1_Utrecht_UU4         -NSDGVWDQIDRVGLLIAIHGTSHYSLLMVLQDGVEASQPHVAVKICHWN--PGNI---S
DQ848678_FCoV1_C1Je_Kitten_FIP     -DSDGVWDQQDKVGLLIAIHGNSHYSLLMVLQDGVEDNQPHVAVKICHWK--PSNI---S
EU186072_FCoV1_Black               -NSDGLWDARDRVGLLIAIYGTSYYSLLMVLQDKVEENQPHVAVKICHWK--PGNI---S
AB088222_FCoV1_UCD1                -NSDGVWDTRDRVGLLIAIHGESHYSLLMVLQDNVEENQPHVAVKICHWQ--LGNI---S
                                      :  *   ::*** : ::* : *.* *:  : ..     .:****.*      .

KP849472.1_CCoV1_Italy-23/03       GGNRYNLHIGDTSDCVFNHRFALDSKLITTDIYGFQWTETYINIYLGGTISRVWIDNTWD
AY307020.1_CCoV1_Elmo/02           ESNRWDLSQGTVPDCKFNHQFALDTKLITSDFYGFQWTNTYVNIYLGGTVSRVWIENTWD
AAB47503_FCoV1_KU-2                SYHAFSVNLGDGGQCVFNQRFSLDTVLTTNDFYGFQWTDTYVDIYLGGTITKVWVDNDWS
KX722530_FCoV1_Dutch_cat2          SYHQFNVEFGDGGQCVFNKRFSLDTVLTTNDFYGFQWTDTYVDIYLGGTITKVWIANDWS
KX722529_FCoV1_Belgium_UG-FH8      SYHQFGVDLGDSGQCVFNRRFSLDTKLTANDFYGFQWTDTYVDIYLGGTITKVWIDNDWS
KP143512_FCoV1_UK26M               SIHQFSVNLGDGGQCVFNQRFSLDTILTTNDFYGFQWTNNYVNIYLGGTITKVWVENDWS
MG893511_FCoV1_Germany_Felix       SYHSFTIDFGDGGQCVFNQRFSLDTKLTTNDFYGFQWTNTYVDIYLGGTITKVWVANDWS
KY566209_FCoV1_China10             IYHKWSASLGDGGQCVFNRRFSLDTVLTANDFYGFQWTDTYVDIYLSGTVTKVWIENDWN
FJ938054_FCoV1_Utrecht_UU4         TYHQFDVNLGDGGQCVFNQRFSLDTVLTANDFYGFQWTDTYVDIYLGGTITKVWVVNDWS
DQ848678_FCoV1_C1Je_Kitten_FIP     SYHQFNVNLGDVGYCVFNRRFSLDTVLTTNDFYGFQWTDTYVDIYLGGTVTKVWIGNDWS
EU186072_FCoV1_Black               SYHQFSVNLEDGGQCVFNQRF--DTVLTANDFYGFQWTDTYVDIYLGGTITKVWVNNDWS
AB088222_FCoV1_UCD1                SYHQFNLNLGDGGQCVFNQRFSLDTVLTANDFYGFQWTDTYVDIYLGGTITKVWVDNDWS
                                     . :          * **::*  *: * :.*:******:.*::***.**:::**: * *.
```

```
KP849472.1_CCoV1_Italy-23/03      KVDVKISSNWNAKTNYGYYVQFVNKTTYYTYNNTGGTGVNQLQLGLCEDDFCAGYVTNVF
AY307020.1_CCoV1_Elmo/02          KVQVKMSPFWSS-LTYGYYFQLVNRTTYYIYNNTGGDDVEQLQLIECKNEFCAGFSKNVF
AAB47503_FCoV1_KU-2               IVEASISYHWNR-INYGYYMQFVNRTTYYAYNNTGGANYTQLQLSECHTDYCAGYAKNVF
KX722530_FCoV1_Dutch_cat2         VVESSISYHWNQ-LNYGYYIQFVNRTTFYIYNNTGGSNYTHLRLSECHGAYCAGYAKNVF
KX722529_FCoV1_Belgium_UG-FH8     VVESSISYYWST-YNKGYYMQFVNRTTYYAYNSTGGSNYTHLQLSECHSEYCAGYAKNVF
KP143512_FCoV1_UK26M              VVESSISYHWSQ-INYGYYMQFVNRTTYYVYNNTGGANYTHLQLEECHSEYCAGYAKNVF
MG893511_FCoV1_Germany_Felix      VVEASISYYWNR-INYGYYIQFVNRTTYYTYNNTGGMNYTHLQLNECTSDYCAGYAKNVF
KY566209_FCoV1_China10            VVEASISYHWNR-INYGYYIQFVNRTTYYAYNSTAGSNYTHLQLKECHSDYCAGYAKNVF
FJ938054_FCoV1_Utrecht_UU4        VVEASISSHWNA-LNYGYYIQFVNRTTYYAYNSTGGSNYTHLQLTECHTDYCAGYAKNVF
DQ848678_FCoV1_C1Je_Kitten_FIP    IVEASISYHWNQ-LNYGYYLQFVNRTTYYTYNSTAGSNYTHLQLAECTSEYCAGYVKNVF
EU186072_FCoV1_Black              SVEASISYHWNQ-INYGYYIQFVNRTTYYTYNNTGGSNYTHLQLAECHSEYCAGYAKNVF
AB088222_FCoV1_UCD1               VVEASISYHWSR-LNYGYYMQFVNRTTYYIYNNTGGSNYTHLHLQECHSDYCAGYAKNVF
                                   *: .:*  *.    . ***.*:**:**:* **.*.*    :*:*  *    :***: .***

KP849472.1_CCoV1_Italy-23/03      TPVNGKIPGDFSFNNWFLLSDKSTIVQGRVVSSQPVYYVECLRPVPSWSNNSAIVSFDNDT
AY307020.1_CCoV1_Elmo/02          TPVNGKIPEDFSFNNWFLLSDKSTIVQGRVVRSQPVYYVECLRPVPSWSNNSAIVSFDNDT
AAB47503_FCoV1_KU-2               VPIDGKIPEDFSFSNWFLLSDKSTLVQGRVLSSQPVFVQCLRPVPSWSNNTAVVHFKNDA
KX722530_FCoV1_Dutch_cat2         VPIDGKIPESFSFSNWFLLSDKSTLVQGRILSKQPVFVQCLRPVPLWSNNSAVVYFKNDA
KX722529_FCoV1_Belgium_UG-FH8     VPIEGKIPESFSFSNWFLLSDKSTLVQGRVLSKQPVFVQCLRPVPSWSNNTAVVLFTNDS
KP143512_FCoV1_UK26M              VPIDGKIPDGFSFSNWFLLSDKSTLVQGRVLSKQPVFVQCLRAVPSWSNNSAVVHFSNDD
MG893511_FCoV1_Germany_Felix      VPVDGKIPESFSFSNWFLLSDKSTLVQGRVLSKQPVFVQCLRPVPSWSNSSAVVYFTNDS
KY566209_FCoV1_China10            VPIDGKIPESFSFSNWFLLSDKSTLVQGRVLSRQPVFVQCLRPVPTWSNNSAVVFFKNDA
FJ938054_FCoV1_Utrecht_UU4        VPIDGKIPEGFSFSNWFLLTDKSTLVQGRVLSSQPVFVQCLRPVPTWSNNTAVVHFKNDV
DQ848678_FCoV1_C1Je_Kitten_FIP    VPIDGKIPESFSFSNWFLLSDKSTLVQGRVLSKQPVFVHCLRPVPSWSNNSAVVHFSDDP
EU186072_FCoV1_Black              VPIDGKIPESFSFSNWFLLSDKSTLVQGRVLSRQPVLVQCLRPVPTWSNNSAVVFFSNDL
AB088222_FCoV1_UCD1               VPIDGKIPESFSFNNWFLLSDKSTLVQGRVLSSQPVFVQCLRPVPTLSNNTAVVHFNNDV
                                  .*::****   ***.*****:****:****::   *** *.*** **   **.:*:* * :*

KP849472.1_CCoV1_Italy-23/03      FCPQASADVLRFNLNFSGEVFHPNATADDQIIFTFEDNSTAEISCYNGANVTDPTIGFNT
AY307020.1_CCoV1_Elmo/02          FCPQASADVLRFNLNFSGEVFHPNATADDQIIFTFEDNSTAEISCYNGANVTDPTVGFNT
AAB47503_FCoV1_KU-2               FCPNVTADVLRFNLNFSDTDVYTDSTNDEQLFFTFEDNTTASIACYSSANVTDFQPANNS
KX722530_FCoV1_Dutch_cat2         FCHNVTTDVLRFNLNFSDTDIYIESTKDDQLHFTFEDNTTASIACYSSANVTDFQPANNS
KX722529_FCoV1_Belgium_UG-FH8     YCPNVTADVLRFNLNFSDTDVYTESTNDDQLYFTFEDNTTASIACYSSANVTDFQPANNS
KP143512_FCoV1_UK26M              FCPNVTAEVLRFNLNFSDTDVYVASNSDDRLYFTFEDNTTAGVACYSSANITDYKPNTNA
MG893511_FCoV1_Germany_Felix      FCPNVTADVLRFNLNFSDTDVYTNSNNDEQLYFTFEDNTNASIACYSSANVTDFQPANNI
KY566209_FCoV1_China10            FCPNVTADVLRFNLNFSDTDVYTESTNDDQLYFTFEDNTTASIACYSSANVTDFQPANNS
FJ938054_FCoV1_Utrecht_UU4        FCPNVTADVLRFNLNFSDTDVYTDSTTDDQLHFTFEDNTTASITCYSSANVTDNQPASGS
DQ848678_FCoV1_C1Je_Kitten_FIP    FCPGVTAEVLRFNLIFSDTAVYTESTSDDQLHFTFEDNSTASIACYSSDNVTDNQPVDNT
EU186072_FCoV1_Black              YCPGVTAEVLRFNLNFSDSDVYTDLTNDDQFYFTFEDNTTASIACYSSANVTDFQPANSS
AB088222_FCoV1_UCD1               FCPNVTADVLRFNLNFSDTDVYTAST-NDQLYFTFEDNTTASIACYSSANVTDNQPANNS
                                  :*  .:::****** **   .:   . :::: ******:.* ::**.. *:**
                                                                                         NTD

KP849472.1_CCoV1_Italy-23/03      TSQIPFGGTADPYLCFVNFSTAALDNQFLGILPPTIREMAFGRDGSIFINGYKYFNMPPI
AY307020.1_CCoV1_Elmo/02          TSQIPFGSTDDPYLCFANFSTAALDNQFLGILPPTVREMAFGRDGSIFINGYKYFNMPPI
AAB47503_FCoV1_KU-2               VSHIPFGKT--AHFCFANFSHSIVSRQFLGILPPTVREFAFGRDGSIFVNGYKYFSLPAI
KX722530_FCoV1_Dutch_cat2         VSHIPFGKT--AYFCFATFSHSVVSRQFLGILPPIVREFAFGRDGSIFVNGYKYFSLPPI
KX722529_FCoV1_Belgium_UG-FH8     ASHIPFGKTMHSYFCFANFSHSVVSRQFLGILPPTVREFAFGRDGSVFVNGYKYFSLPPI
KP143512_FCoV1_UK26M              SAQIPFGKTTHSYFCFANFSSSHV-TQFLGILPPTVREFAFGKDGSIFVNGYKYFSFPPI
MG893511_FCoV1_Germany_Felix      NSHIPFGTTAQSYFCFANFSNAAVARQFLGILPPTVREFAFGRDGSVFVNGYKYFSLPPI
KY566209_FCoV1_China10            VSHVPFGKTEHSYFCFANFSHAVVSRQFLGILPPTVREFAFGRDGSIFVNGYKYFSLPPI
FJ938054_FCoV1_Utrecht_UU4        ISHTPFVS--NSYLCFANFSHSSVSRQFLGILPPTVREFAFGRDGSIFVNGYKYFSLQPI
DQ848678_FCoV1_C1Je_Kitten_FIP    LSDIPFGKITHSYLCFANFSNAPVSMQFLGILPPTVREFAFGRDGSIFVNGYKYFSFPPI   RBD
EU186072_FCoV1_Black              VSHIPLGKTRHSYFCFANFSHSVVSRQVLGILPPTVREFAFGRDGSVFVNGYKYFSLPPI   55T
AB088222_FCoV1_UCD1               TSHIPFGKTMHSYFCFANFSNS-MVRQFLGILPPTVREFAFGRDGSIFVNGYKYFSLPPI   27I
                                   :. *:       ::**..** : :   *.****** :**:***:***:*:******.:  *   42S
                                        Proposed RBD / S1

KP849472.1_CCoV1_Italy-23/03      KSINFAISAVEYYGFWTIAYTKYTDVMVDVNGTSITRLFYCDSPLNRIKCQQMKHELPDG
AY307020.1_CCoV1_Elmo/02          KSINFAISAVEYYGFWTIAYTKYTDVMVDVNGTSITRLFYWDSPINRIKGQQKKHDLPDG
AAB47503_FCoV1_KU-2               RSVNFSISSVEEYGFWTIAYTNYTDVMVDVNGTAITRLFYCDSPLNRIKCQQLKHELPDG
KX722530_FCoV1_Dutch_cat2         KSVNFSISSVEQYGFWTIAYTNYTDVMVDVNGTGITRLFYCDSPLNRIKCQQLKYELPDG
KX722529_FCoV1_Belgium_UG-FH8     KSVNFSISSVEQYGFWTIAYTNYTDVMVDVNGTVITRLFYCDSPLNRIKCQQLKHELPDG
KP143512_FCoV1_UK26M              KSVNFSISSVENFGFWTIAYTNYTDVMVDVNGTGITRLFYCDSPINRIKCQQLKYELPDG
MG893511_FCoV1_Germany_Felix      KSVNFSISSIQRYGFWTIAYTNYTDVMVDVDGTGITRLFYCDSPINRIKCQQLKHELPDG
KY566209_FCoV1_China10            KSVNFSISSVEQYGFWTIAYTNYTDVMVDVNGTSITRLFYCDSPLNRIKCQQLKHELPDG
FJ938054_FCoV1_Utrecht_UU4        KSVNFSISSVENYGFWTIAYTNYTDVMVDVNGTVITRLFYCDSPLNRIKCQQLKHELPDG
DQ848678_FCoV1_C1Je_Kitten_FIP    KSVNFSISSVENYGSWTIAYTNYTDVMVDVNGTGITRLFYCDSPLNRIKCQQLKYELPDG
EU186072_FCoV1_Black              KSVNFSISSVEQYGFWTIAYTNYTDVMVDVNGTGISRLFYCDSPLNRIKCQQLKHVLPDG   60T
AB088222_FCoV1_UCD1               KSVNFSIGSVEQHGFWTIAYTNYTDVMVDINGTGITRLFYCDSPINRIKCQQLKHELPDG   39I
                                  :*:**:*.::: .* ******:*******::** *:**** ***:**** ** *: ****   53S
```

```
KP849472.1_CCoV1_Italy-23/03        FYSANSLVKKNLPKTFVTTPQFYNGMNVTLHVILNDTSAGTKVVGFERYELEQIADILFE
AY307020.1_CCoV1_Elmo/02            FYSANSLVKKDLPKTFVTTPQFYNWMNVTLHVILNDTEADYKVVGFERYELEKIADIHFE
AAB47503_FCoV1_KU-2                 FYSASMLVKKDLPKTFVTMPQFYHWMNVTLHVVLNDTEKKYDIILAKAPELAALADVHFE
KX722530_FCoV1_Dutch_cat2           FYSASMLVKKDLPKTFVTMPQFYNWMNVTLHVVLNDTEKKPDTILAKAPELASLADIHFE
KX722529_FCoV1_Belgium_UG-FH8       FYSASMLVKKDLPKTFVTMPQFYNWMNVTLHVVLNDTEKRQDIILAKASELASLADIHFE
KP143512_FCoV1_UK26M                FYSASMLVKKDLPKTFVTMPQFYNWMNVTLHVVLNDTEKKADIILAKASELASLADVHFE
MG893511_FCoV1_Germany_Felix        FYSASTLVKKDLPKTFVTMPQFYNWMNVTLHVVLNDTQKGANIILSKAAELASLADIHFE
KY566209_FCoV1_China10              FYSASTLVKKDLPKTFVTMPQFYNWMNVTLHVVLNDTEKKEDIILAKAPELLSLADIHFE
FJ938054_FCoV1_Utrecht_UU4          FYSASMLVKKDLPKTFVTMPQFYNWMNVTLHVVLNDIEKKADIILAGAPELASLADIHFE
DQ848678_FCoV1_C1Je_Kitten_FIP      FYSASALVKKDLPKTFVIMPQFYNWMNVTLHVVLNDTAKGADIILAKAPELASLADVHFE RBD
EU186072_FCoV1_Black                FYSASMLVKNDLPKTFVTMPQFYNWMHITLHVVLNDTAKKEDIILAKAPELASLADVHFE 60T
AB088222_FCoV1_UCD1                 FYAASMLVKKDLPKTFVTMPQFYNWMTVTLHVVLNDTEKKADIILAKAPELASLADIHFE 30I
                                    **:*. ***::******  ****. * :****:***      . :     **  :**: ** 41S

KP849472.1_CCoV1_Italy-23/03        IPQNQDGEVKNVTNFCVQTRQLAFFFKYTSLQGLYSISNTVWLKNYDCPFSPQQFNNYLQ
AY307020.1_CCoV1_Elmo/02            IPQDKNGGVKNVTNFCVQTRQLAFFFKYTSLQGLYSISNTVWLKNYDCPFSPQQFNNYLQ
AAB47503_FCoV1_KU-2                 IAQA-NGSVTNVTSLCVQARQLALFYKYTSLQGLYTYSNLVELQNYDCPFSPQQFNNYLQ
KX722530_FCoV1_Dutch_cat2           VAQS-NGSVTNVTSICVQTRQLALFYKYTSLQGLYTYSNLVELQNYDCPFSPQQFNNYLQ
KX722529_FCoV1_Belgium_UG-FH8       IAQA-NGSVTNVTSLCIQARQLALFYKYTSLQGLYTYSNLVEVQNHDCPFSPQQFNNYLQ
KP143512_FCoV1_UK26M                IAQA-NGSVVNATSLCVQTRQLALFYKYTSLQGLYTYSNLVELQNYDCPFSPHQFNNYLQ
MG893511_FCoV1_Germany_Felix        IDRA-DGSVTNVTSLCVQTRQLALFYKYTSLQGLYTYSNLVELQNYDCPFSPQQFNNYLQ
KY566209_FCoV1_China10              IAQA-NGSVTNVTSVCVQARQVALFYKYTSLQGLYTYSNLVELQNYDCPFSPQQFNNYLQ
FJ938054_FCoV1_Utrecht_UU4          IAQA-NGSVVNVTSVCVQARQLALFYKYTSLQGLYTYSNLVQLQNYDCPFSPQQFNNYLQ
DQ848678_FCoV1_C1Je_Kitten_FIP      IAQA-NGSVANLTSLCVQTRQLALFYKYTSLQGLYTYSNLIELQNYDCPFSPQQFNNYLQ RBD
EU186072_FCoV1_Black                ILQA-NGSVVNVTSLCVQAKQLALFYKYTSLQGLYTYSNLVELQNYDCPFSPQHFNNYLQ 60T
AB088222_FCoV1_UCD1                 IVQA-NGSVANVTSLCVQTRQIALFYKYTSLQSLYTYSNLVELQNYDCPFSPQQFNNYLQ 32I
                                    : :  :* * * *..*:*::*:*:*:******.**: ** : ::*:******::****** 51S

KP849472.1_CCoV1_Italy-23/03        FETLCFDTNPSVAGCKWAVMHSPIWNTQFATITVTYKEGAQITTMPK-PQLGFRDVSFKV
AY307020.1_CCoV1_Elmo/02            FETLCFDTNPSVAGCKWAIMHSPIWNTQFATITVTYKDGAQITTMPK-PQLGFRDISFKV
AAB47503_FCoV1_KU-2                 FETLCFDVNPAVAGCKWSLVHDVQWRTQFATITVSYKHGSMITTHAKGHSWGFQDTSVLV
KX722530_FCoV1_Dutch_cat2           FETLCFDVNPAVAGCKWSLVHDTRWRTQFATITVSYKEGSMITTMPS-SQLGFQDISVLV
KX722529_FCoV1_Belgium_UG-FH8       FETLCFDVNPAVAGCKWSLVHDNIWRTQFATITVSYKEGAMITTMPK-AQLGFQDISVLV
KP143512_FCoV1_UK26M                FETLCFDTSPAVAGCKWSLVHDVRWRTQFATITISYKDGAKITTMPK-AKLGFQDISNIV
MG893511_FCoV1_Germany_Felix        FETLCFDVNPSVAGCKWSLLHDSIWHTQFATITVSYKEGARITTMPK-AQIGFQDISNLV
KY566209_FCoV1_China10              FETLCFDVSPAVAGCKWSLVHDNKWRTQFATITVSYKEGSMITTMPK-PQLGFQDISNLV
FJ938054_FCoV1_Utrecht_UU4          FETLCFDVSPAVAGCKWSLVHDVKWRTQFATITVSYKDGAMITTMPK-AQLGFQDISNIV
DQ848678_FCoV1_C1Je_Kitten_FIP      FETLCFDVSPVVAGCKWALVHDYNWRTQFATITVSYKEGAMITTMPK-AQLGFQDISKLV RBD
EU186072_FCoV1_Black                FETLCFDVNPAVAGCKWSLVHDVNWRTQFATITVSYKDGAMITTMPR-AQLGFKDISNLV 46T
AB088222_FCoV1_UCD1                 FETLCLDTNPAVAGCKWALVHDVKWRTQFATITVSYKDGAMITTMPK-AQLGFQDISNLV 28I
                                    *****:*..* ******:::*.   *.*******::**.*: ***      . **:* *  * 40S
```

**Proposed RBD**

RBD Total (T): 281 aa
Identity (I): 156 / 281 = 55.5%
Similarity(S): 227 / 281 = 80.8%

```
KP849472.1_CCoV1_Italy-23/03        EGECTDYNIYGFQGTGIIRKTESKLVAGLFYTSASGDLLAFKNSTTGEIFTVVPCDLTAQ
AY307020.1_CCoV1_Elmo/02            EGECTDYNIYGFQGTGIIRKTESKLVAGLFYTSASGDLLAFKNSTTGEIFTVIPCDLTAQ
AAB47503_FCoV1_KU-2                 KDECTDYNIYGFQGTGIIRNTTSRLVAGLYYTSISGDLLAFKNSTTGEIFTVVPCDLTAQ
KX722530_FCoV1_Dutch_cat2           KDECTDYNIYGFQGTGVIRNTTSRLVAGLYYTSISGDLLAFKNSTTGEIFTVVPCDLTAQ
KX722529_FCoV1_Belgium_UG-FH8       KDECTDYNIYGFQGTGIIRNTTSRLVAGLYYTSISGDLLAFKNSTTGEIFTVVPCDLTAQ
KP143512_FCoV1_UK26M                KDECTDYNIYGFQGTGIIRNTTTRIVAGLYYTSIIGDLLAFKNSTTGEIFTVVPCGLTAQ
MG893511_FCoV1_Germany_Felix        KDECTDYNIYGFHGTGIIRPTTSRLVAGLYYTSTSGDLLAFKNSTTGEIFTVTPCDLTAQ
KY566209_FCoV1_China10              KDECTDYNIYGFQGTGIIRNTTSRLVAGLYYTSISGDLLAFKNSTTGEIFTVVPCDLTAQ
FJ938054_FCoV1_Utrecht_UU4          KDECTDYNIYGFQGTGIIRSTTSRLVAGLYYTSASGDLLGFKISTTGEIFTVVPCDLTAQ
DQ848678_FCoV1_C1Je_Kitten_FIP      KDECTDYNIYGFQGTGIIRKTTSRLVAGLYYTSISGDLLAFKNSTTGEIFTVVPCDLTAQ
EU186072_FCoV1_Black                KDECTDYNIYGFQGTGIIRSTTSRLVAGLYYTSISGDLLAFKNSTTGEIFTVVPCDLTVL
AB088222_FCoV1_UCD1                 KDECTDYNIYGFHGTGIIRNTTSRLVAGLYYTSISGDLLAFKNSTTGEIFTVVPCDLTAQ
                                    : **********:***:** * :::****:***  ****.** ********* ** **.
```

**CTD / S1**

```
KP849472.1_CCoV1_Italy-23/03      AAVINDEIVGAITSINQTDLFEFVNHTHTVRRVRRAVQTGTTITAYSMPQFYYITKWNND
AY307020.1_CCoV1_Elmo/02          AAVINDEIVGAITSINQTDLFEFVNHTHTVRRARRAVQTGTTITAYSMPQFYYITKWNND
AAB47503_FCoV1_KU-2               VAVINDEIVGAITAVNQTDLFEFVNNTQARRSRS---STPNFVTSYTMPQFYYITKWNND
KX722530_FCoV1_Dutch_cat2         AAVINDEIVGVITSVNQTDLFEFVNHTSTRRSRI--AAVPQAATTYTMPQFYYITKWNND
KX722529_FCoV1_Belgium_UG-FH8     AAVINDEIVGAITAVNQTDLFEFVNHTQVKRLRR--S-TPETVQTYTMPQFYYITKWNND
KP143512_FCoV1_UK26M              AAVINDEIVGVITAVNQTDLFEFVNHTQARGARR--STGSQTVQTYTMPQFYYITKWNND
MG893511_FCoV1_Germany_Felix      AAVINDEIVGAITAVNQTDLFEFVNHTSHRRSRR--E--VPTVQTYTMPQFYYITKWNND
KY566209_FCoV1_China10            AAVINDEIVGAITAVNQTDLFEFVNHTHTRRSRR--S-PTEAVKTYTMPQFYYITKWNND
FJ938054_FCoV1_Utrecht_UU4        AAVINDEIVGAITATNQTDLFEFVNHTWSRSARG--SS-PSTVNTYTMPQFYYITKWNNG
DQ848678_FCoV1_C1Je_Kitten_FIP    AAVINDEIVGVITAVNQTDLFEFVNHTQPRQSRR--SANPTTVQTYTMSQFYYITKWNND
EU186072_FCoV1_Black              AAVINDEIVGAITSVNQTDLFEFFNHTQAKRSRR--PT-SHSVTTYNMPQFYYITRWNND
AB088222_FCoV1_UCD1               AAVINDEIVGVITAVNQTDLFEFVNHTHSRRSRG--ST-STSVTTYTMPQFYYITKWNND
                                  .*********.**: ********.*.*            :*.* ******:***
```

S1/S2

CTD

```
KP849472.1_CCoV1_Italy-23/03      TSENCTSVITYSSFAICNTGEIKFVNVTKVEAVEDGIGTVKPISTGNITIPKNFTVAVQA
AY307020.1_CCoV1_Elmo/02          TSENCTSVITYSSFAICNTGEIKYVNVTKVEAVEDGIGTIKPISTGNITIPKNFTVAVQA
AAB47503_FCoV1_KU-2               TSSNCTSAITYSSFAICNTGEIKYVNVTHVEIVDDSIGVIKPVSTGNISIPKNFTVAVQA
KX722530_FCoV1_Dutch_cat2         TSTNCTSVITYSSFAICNTGEIKYVNVTHVEVV-----VVKPVSTGNITIPKNFTVAVQA
KX722529_FCoV1_Belgium_UG-FH8     TSTNCTSVITYSSFAICNTGEIKYVNVTKVEIVDDSIGVIKPVSTGNISIPKNFTVAVQA
KP143512_FCoV1_UK26M              SSTNCTSVITYSSFAICNTGEIKYVNVTHVEIVDDSIGVIKPISTGNIFIPKNFTVAVQA
MG893511_FCoV1_Germany_Felix      TSTNCTSVITYSSFAICNTGEIKYVNVTKVEVVDDSIGVIKPISSGNISIPKNFTVAVQA
KY566209_FCoV1_China10            TSTNCTSVITYSSFAICNTGEIKYVNVTHVETVDDNIGVIRPISTGNISIPKNFTVAVQA
FJ938054_FCoV1_Utrecht_UU4        TSSNCTSVITYSSFAICNTGEIKYVNVTHVEIVDDSVGVIKPVSTGNITIPKNFTVAVQA
DQ848678_FCoV1_C1Je_Kitten_FIP    TSTNCTSFITYSSFAICNTGEIKYVNVTHVETVDDSIGVIKPISTGNISIPKNFTVAVQA
EU186072_FCoV1_Black              TSTNCTSVITYSSFAICNTGEIKYVNVTHVETVDD-IGVIKPISTGNILIPKNFTVAVQA
AB088222_FCoV1_UCD1               TSTNCTSVITYSSFAICNTGEIKYVK-------DDSIGVIKPISTGNISIPKNFTVAVQA
                                  :* **** ***************:*:           .::*:*:*** ***********
```

```
KP849472.1_CCoV1_Italy-23/03      EYIQIQVKPVVVDCAKYVCNGNNHCNRLLTQYTSACQNIENALNLGARLESLMLNEMITV
AY307020.1_CCoV1_Elmo/02          EYIQIQVKPVVVDCAKYVCNGNNHCNKLLTQYTSACQNIENALNLGARLESLMLNEMITV
AAB47503_FCoV1_KU-2               EYIQIQVKPVVVDCATYVCNGNTHCLKLLTQYTSACQTIENALNLGARLESLMLNDMITV
KX722530_FCoV1_Dutch_cat2         EYIQIQVKPVAVDCAKYVCNGNRHCLNLLTQYTSACQTIENALNLGARLESLMLNDMITV
KX722529_FCoV1_Belgium_UG-FH8     EYIQIQVKPVVVDCAKYVCNGNRHCLNLLTQYTSACQTIENALNLGARLESLMLNDMVTV
KP143512_FCoV1_UK26M              EYIQIQVKPVVVDCAKYVCNGNSHCLTLLAQYTSACQTIENALNFGARLESLMLNDMITV
MG893511_FCoV1_Germany_Felix      EYIQIQVKPVVVDCAKYVCNGNKHCLNLLTQYTSACQTIENALNLGARLESLMLNDMITV
KY566209_FCoV1_China10            EYIQIQVKPVVVDCAKYVCNGNRHCLSLLAQYTSACQTIENALNLGARLESLMLNDMITV
FJ938054_FCoV1_Utrecht_UU4        EYVQIQVKPVAVDCAKYVCNGNRHCLNLLTQYTSACQTIENSLNLGARLESLMLNDMITV
DQ848678_FCoV1_C1Je_Kitten_FIP    EYFQIQVKPVVVDCAKYVCNGNRHCLSLLAQYTSACHTIENALNLGARLESQMLKDMITV
EU186072_FCoV1_Black              EYIQIQVKPVVVDCAKYVCNGNTHCRSLLTQYTSACQTIENALNLGARLESLMLNDMITV
AB088222_FCoV1_UCD1               EYIQIQVKPVVLDCAKYVCNGNSHCLSLLTQYTSACQTIENALNLGARLESLMLNDMITV
                                  **.*******.:***.****** **  **:******:.***:**:****** **::*:**
```

```
KP849472.1_CCoV1_Italy-23/03      SDSSLELATIEKFNTTVVGGEKLGGIYFDGLKDVLPAQPGGRSAIEDLLFNKVVTSGLGT
AY307020.1_CCoV1_Elmo/02          SDSSLELATIEKFNTTVVGGEKLGGIYFDGLKDVLPAQPGGRSAIEDLLFNKVVTSGLGT
AAB47503_FCoV1_KU-2               SDRGLELATVERFNATALGGEKLGGLYFDGLSSLLPPKIGKRSAVEDLLFNKVVTSGLGT
KX722530_FCoV1_Dutch_cat2         SYRSLELATVEKFNTTVVGGEKLGGFYFDGLRALLPPTIGKRSAVEDLLFNKVVTSGLGT
KX722529_FCoV1_Belgium_UG-FH8     SDRSLELATVEKFNTTLVGGEKLGGFYFDGLSSLLPPKIGKRSAVEDLLFNKVVTSGLGT
KP143512_FCoV1_UK26M              SDRSLELATVEKFNTTVLGAEKLGGFYFDGLRELLPPTIGKRSAIEDLLFNKVVTSGLGT
MG893511_FCoV1_Germany_Felix      SDRSLELATVEKFNSTVLSGEKLGGLYFDGLSSFLPPRIGKRSAIEDLLFNKVVTSGLGT
KY566209_FCoV1_China10            SDRSLELANVEKFNTTLLGGEKLGGFYFDGLRELLPPTIGKRSAVEDLLFNKVVTSGLGT
FJ938054_FCoV1_Utrecht_UU4        SDRSLEFATVDKFNTTALGGEKLGGLYFDGLSSLLPPRVGMRSAVEDLLFNKVVTSGLGT
DQ848678_FCoV1_C1Je_Kitten_FIP    SDRSIQLATVEKFNNTLLGGEKLGGHYFDGLKSLLPPRIGKRSAVEDLLFDKVVTSGLGT
EU186072_FCoV1_Black              SDRSLELATIEKFNATATGGVKLGGLYFDGLSSLLPPKIGVRSAVEDLLFNKVVTSGLGT
AB088222_FCoV1_UCD1               SSRSLELATVERFNATAPGGEKLGGLYFDGLSSLLPPRVGQRSAVEDLLFNKVVTSGLGT
                                  *  .:::*.:::** *  .. **** *****   .**    * ***:*****:*********
```

S2’

```
KP849472.1_CCoV1_Italy-23/03       VDEDYKKCSAGTDIADLVCAQYYNGVMVLPGVVDDNKMAMYTASLIGGMAMGSITSAVAI
AY307020.1_CCoV1_Elmo/02           VDEDYKKCSAGTDIADLVCAQYYNGVMVLPGVVDDNKMAMYTASLIGGMAMGSITSAVAI
AAB47503_FCoV1_KU-2                VDDDYKKCSSGTDVADLVCAQYYNGIMVLPGVVDGNKMSMYTASLIGGMALGSITSAVAV
KX722530_FCoV1_Dutch_cat2          VDDDYKKCSSGTDVADLVCAQYYNGIMVLPGVVDGNKMAMYTASLIGGMALGSITSAVAV
KX722529_FCoV1_Belgium_UG-FH8      VDDDYKKCSAGTDVADLVCAQYYNGIMVLPGVVDDNKMAMYTASLIGGMAMGSITSAVAV
KP143512_FCoV1_UK26M               VDDDYKKCSAGTDVADLVCAQYYNGIMVLPGVVDQNKMSMYTASLIGGMALGSITSAVAV
MG893511_FCoV1_Germany_Felix       VDDDYKKCSSGIDVADLVCAQYYNGIMVLPGVVDDNKMAMYTASLIGGMAMGSITSAVAV
KY566209_FCoV1_China10             VDDDYKKCSSGTDVADLVCAQYYNGIMVLPGVVDQNKMAMYTASLIGGMAMGSITSAVAV
FJ938054_FCoV1_Utrecht_UU4         VDDDYKKCSAGTDVADLVCAQYYNGIMVLPGVVDYNKMAMYTASLIGGMALGSITSAVAV
DQ848678_FCoV1_C1Je_Kitten_FIP     VDDDYKKCSAGTDVADLVCAQYYNGIMVLPGVVDQNKMAMYTASLIGGMALGSITSAVAV
EU186072_FCoV1_Black               VDDDYKKCSAGTDVADLVCAQYYNGIMVLPGIVDGNKMAMYTASLIGGMALGSITSAVAV
AB088222_FCoV1_UCD1                VDDDYKKCSAGTDVADLVCAQYYNGIMVLPGVVDGNKMAMYTASLIGGMALGSITSAVAV
                                   **:******:* *:***********:*****:** ***:***********:********:

KP849472.1_CCoV1_Italy-23/03       PFAMQVQARLNYVALQTDVLQENQKILANAFNNAMGNITLALGQVSDAISTVSEGFNTMA
AY307020.1_CCoV1_Elmo/02           PFAMQVQARLNYVALQTDVLQENQKILANAFNNAIGNITLALGQVSDAISTVSEGFNTMA
AAB47503_FCoV1_KU-2                PFAMQVQARLNYVALQTDVLQENQKILANAFNNAIGNITLALGKVSNAITTTSDGFNSMA
KX722530_FCoV1_Dutch_cat2          PFAMQVQARLNYVALQTDVLQENQKILANAFNNAIGNITLALGKVSNAITTTSYGFNSMA
KX722529_FCoV1_Belgium_UG-FH8      PFAMQVQARLNYVALQTDVLQENQKILANAFNNAIGNITLALEKVSNAITTISDGFNSMA
KP143512_FCoV1_UK26M               PFAMQVQARLNYVALQTDVLQENQKILANAFNNAIGNITLALGKVSNAITTTSDGFNTMA
MG893511_FCoV1_Germany_Felix       PFAMQVQARLNYVALQTDVLQENQKILANAFNNAIGNITLALGKVSNAITTISDGFNTMA
KY566209_FCoV1_China10             PFAMQVQARLNYVALQTDVLQENQKILANAFNNAIGNITLALGKVSNAITTISDGFNSMA
FJ938054_FCoV1_Utrecht_UU4         PFSMQVQARLNYVALQTDVLQENQKILANAFNNAIGNITLALGKVSNAITTVSDGFNSMA
DQ848678_FCoV1_C1Je_Kitten_FIP     PFAMQVQARLNYVALQTDVLQENQKILANAFNNAIGNITLALGKVSNAITTISDGFNTMA
EU186072_FCoV1_Black               PFSMQVQARLNYVALQTDVLQENQKILANAFNNAIGNITLALGKVSNAVTTISDGFNTMA
AB088222_FCoV1_UCD1                PFAMQVQARLNYVALQTDVLQENQKILANAFNNAIGNITLALGKVSNAITTISDGFNTMA
                                   **:*******************************:******* :**:*::* * ***:**

KP849472.1_CCoV1_Italy-23/03       SALTKIQSVVNQQGNALSQLTQQLQNNFQAISSSIAEIYNRLEKVEADAQVDRLITGRLA
AY307020.1_CCoV1_Elmo/02           SALTKIQSVVNQQGNALSQLTQQLQNNFQAISSSIAEIYNRLEKVEADAQVDRLITGRLA
AAB47503_FCoV1_KU-2                SALTKIQSVVNQQGEALSQLTSQLQKNFQAISSSIAEIYNRLEKVEADAQVDRLITGRLA
KX722530_FCoV1_Dutch_cat2          LALTKIQSVVNQQGEALSQLTSQLQKNFQAISSSIAEIYNRLEKVEADAQVDRLITGRLA
KX722529_FCoV1_Belgium_UG-FH8      SALTKIQSVVNQQGEALAQLTSQLQKNFQAISSSIAEIYNRLEKVEADAQVDRLITGRLA
KP143512_FCoV1_UK26M               SALTKIQSVVNHQGDALSQLTSQLQNNFQAISSSIAEIYNRLEKVEADAQVDRLITGRLA
MG893511_FCoV1_Germany_Felix       SALTKIQSVVNQQGEALSQLTGQLQKNFQAISSSIAEIYSRLEKVEADAQVDRLITGRLA
KY566209_FCoV1_China10             SALTKIQSVVNQQGEALSQLTSQLQKNFQAISSSIAEIYNRLEKVEADAQVDRLITGRLA
FJ938054_FCoV1_Utrecht_UU4         SALTKIQSVVNQQGEALSHLISQLQKNFQAISSSIAEIYNRLEKVEADAQVDRLITGRLA
DQ848678_FCoV1_C1Je_Kitten_FIP     SALTKIQSVVNQQGEALSQLTSQLQKNFQAISSSIAEIYNRLEKVEADAHVDRLITGRLA
EU186072_FCoV1_Black               SALTKIQSVVNQQGEALSQLTSQLQKNFQAISSSIAEIYNRLEKAEADAQVDRLITGRLA
AB088222_FCoV1_UCD1                SALTKIQSVVNQQGEALSLLTSQLRKNFQAISSSIAEIYNRLEKVEADAQVDRLITGRLA
                                    **********:**:**: *  **::*************.****.****:**********

KP849472.1_CCoV1_Italy-23/03       ALNAYVAQTLTQYTEVKASRQLAMEKVNECVKSQSDRYGFCGNGTHLFSLANAAPDGLLF
AY307020.1_CCoV1_Elmo/02           ALNAYVAQTLTQYTEVKASRQLAMEKVNECVKSQSDRYGFCGNGTHLFSLANAAPDGLLF
AAB47503_FCoV1_KU-2                ALNAYVSQTLTQYAEVKASRQIALEKVNECVKSQSNRYGFCGNGTHLFSLVNSAPEGLLF
KX722530_FCoV1_Dutch_cat2          ALNAYVSQTLTQYAEVKASRQLAMEKVNECVKSQSDRYGFCGNGTHLFSLVNSAPDGLLF
KX722529_FCoV1_Belgium_UG-FH8      ALNAYVSQTLTQYAEVKASRQLAMEKVNECVKSQSDRYGFCGNGTHLFSLVNSAPDGLLF
KP143512_FCoV1_UK26M               ALNAYVSQTLTQYSEVKASRQLAMEKVNECVKSQSNRYGFCGNGTHLFSLVNSAPDGLLF
MG893511_FCoV1_Germany_Felix       ALNAYVSQTLTQYAEVKASRQLAMEKVNECVKSQSDRYGFCGNGTHLFSLVNSAPDGLLF
KY566209_FCoV1_China10             ALNAYASQTLTQYAEVKASRQLAMEKVNECVKSQSDRYGFCGNGTHLFSLVNSAPDGLLF
FJ938054_FCoV1_Utrecht_UU4         ALNAYVAQTLTQYAEVKASRQLAMEKVNECVKSQSDRYGFCGNGTHLFSLVNSAPDGLLF
DQ848678_FCoV1_C1Je_Kitten_FIP     ALNAYVAQTLTQYTEVKASRQLAMEKVNECVKSQSNRYGFCGNGTHLFSLVNSAPDGLLF
EU186072_FCoV1_Black               ALNAYVSQTLTQYAEVKASRQLAMEKVNECVKSQSDRYGFCGNGTHLFSLVNSAPDGLLF
AB088222_FCoV1_UCD1                ALNAYVSQTLTQYAEVKASRQLAMEKVNECVKSHSDRYGFCGDGTHLFSLVNSAPDGLLF
                                   *****.:******:*******:*:*********:*:******:*******.*:**:****

KP849472.1_CCoV1_Italy-23/03       LHTVLLPTEWEEVTAWSGICVNDTYAYVLKDFKSSIFSYSGTYMITPRNMFQPRKPQMSD
AY307020.1_CCoV1_Elmo/02           LHTVLLPTEWEEVTAWSGICVNDTYAYVLKDFKSSIFSYNGTYMITPRNMFQPRKPQMSD
AAB47503_FCoV1_KU-2                FHTVLLPTEWEEVTAWSGICVNDTYAYVLKDFDHSIFSYNGTYMVTPRNMFQPRKPQMSD
KX722530_FCoV1_Dutch_cat2          FHTVLLPTEWEEVTAWSGICVNDTYAYVLKDFEYSIFSYNNTYMVTPRNMFQPRKPHMSD
KX722529_FCoV1_Belgium_UG-FH8      FHTVLLPTEWEEVTAWSGICVNDTYAYVLKDFEYSIFSYNNTYMVTPRNMFQPRKPQMSD
KP143512_FCoV1_UK26M               FHTVLLPTEWEVVTAWSGICVNDTYAYVLKDFEYSIFSYNGTYMVTPRNMFQPRKPQMSD
MG893511_FCoV1_Germany_Felix       FHTVLLPTEWEEVTAWSGICVNDTYAYVLKDFEYSIFSYNGTYMVTPRNMFQPRKPQMSD
KY566209_FCoV1_China10             FHTVLLPTEWEEVTAWSGICVNDTYAYVLKDFEYSIFSYNNTYMVTPRNMFQPRKPQMSD
FJ938054_FCoV1_Utrecht_UU4         FHTVLLPTEWEEVTAWSGICVNDTYAYLLKDFDHSIFSYNGTYMVTPRNMFQPRKPQMSD
DQ848678_FCoV1_C1Je_Kitten_FIP     FHTVLLPTEWEEVTAWSGICVNDTYAYVLKDFESSIFSYNNTYMLTPRNMFQPRKPQMSD
EU186072_FCoV1_Black               FHTVLLPTEWEKVTAWSGICVNHTYAYVLKDFDHSIFSYNNTYMVTPRNMFQPRKPQMSD
AB088222_FCoV1_UCD1                FHTVLLPTEWEEVTAWSGICVNDTFAYVLKDFEYFIFSYNNTYMVTPRNMFQPRKPQMSD
                                   :********** **********.*:**:****.  ****. ***:***********:***
```

```
KP849472.1_CCoV1_Italy-23/03       FVQITSCEVTFLNTTYTTFEEIVIDYIDINKTISDMLEQYSPNFTIPDFTEGLEIFNQTK
AY307020.1_CCoV1_Elmo/02           FVQITSCEVTFLNTTYTTFEEIVIDYIDINKTISDMLEQYSPNFTIPDFTEGLEIFNQTK
AAB47503_FCoV1_KU-2                FVQITSCEVTFLNMTYTTFQEIVIDYIDINKTIADMLEQYNPNYTTPELNLLLDIFNQTK
KX722530_FCoV1_Dutch_cat2          FVQITSCEVTFLNTTYTTFQEIVIDYIDINKTISDMLEQYNPNHTIPDLDLQLEIFNQTK
KX722529_FCoV1_Belgium_UG-FH8      FVRITSCEVTFLYTTYTAFQEIVIDYIDINKTISDMLEQYNPNYTTPELDLQLEIFNQTK
KP143512_FCoV1_UK26M               FVQITSCEVTFLNTTYTTFQEIVIDYIDINKTIADMLEQYNPNYTTPELNLQLEIFNQTK
MG893511_FCoV1_Germany_Felix       FVQITSCEVTFLNTTYTTFQEIVIDYIDINKTISDMLEQYNPNYTTPELDLQLDFFNQTK
KY566209_FCoV1_China10             FVQITSCEVTFLNTTYTTFQEIVIDYIDINKTIADMLEQYNPNYTTPELNLQLEIFNQTK
FJ938054_FCoV1_Utrecht_UU4         FVQITSCEVTFLNTTHTTFQEIVIDYIDINKTIADMLEQYHSNYTTPELDLQLEIFNQTK
DQ848678_FCoV1_C1Je_Kitten_FIP     FVQITSCEVTFLNTTYTNFQDIVVDYIDINKTIADMLEQYNPNYTTPDFDLHIEIFNQTK
EU186072_FCoV1_Black               FVQITSCEVTFLNTTYTMFQNIVVDYIDINKTIADMLEQYYSNYTTPELDLQLEIFNQTK
AB088222_FCoV1_UCD1                FVQIMSCEVTFLNTTYTTFQEIVIDYIDINKTIADMLEQYYSNYTTPELDLQLEIFNQTK
                                   **:* *******  *:* *::**:*********:******  *.* *::   :::*****

KP849472.1_CCoV1_Italy-23/03       LNLTAEIDELQVRADNLTVIAHNLQEYIDNLNKTLVDLEWLNRIETYVKWPWYVWLLIGL
AY307020.1_CCoV1_Elmo/02           LNLTAEIDELQVRADNLTVIAHNLQEYIDNLNKTLVDLEWLNRIETYVKWPWYVWLLIGL
AAB47503_FCoV1_KU-2                LNLTAEIDQLEQRADNLTTIAHELQQYIDNLNKTLVDLDWLNRIETYVKWPWYVWLLIGL
KX722530_FCoV1_Dutch_cat2          LNLTAEIDQLEQRADNLTTIAHELQQYIDNLNKTLVDLEWLNRIETYVKWPWYVWLLIGL
KX722529_FCoV1_Belgium_UG-FH8      LNLTAEIDQLEERADNLTVIAHELQQYIDNLNKTLVDLEWLNRVETYVKWPWYVWLLIGL
KP143512_FCoV1_UK26M               LNLTAEIDQLEQRADNLTNIAHQLQQYIDNLNKTLVDLEWLNRVETYVKWPWYVWLLIGL
MG893511_FCoV1_Germany_Felix       LNLTAEIDQLEQRADNLTIIAHELQQYIDNLNKTLVDLEWLNRVETYVKWPWYVWLLIGL
KY566209_FCoV1_China10             LNLTAEIDQLEQRADNLTTIAHELQQYIDNLNKTLVDLEWLNRVETYVKWPWYVWLLIGL
FJ938054_FCoV1_Utrecht_UU4         LNLTAEIDQLEQRADNLTNIAHELQQYIDNLNKTIVDLEWLNRIETYVKWPWYVWLLIGL
DQ848678_FCoV1_C1Je_Kitten_FIP     LNLTAEIDQLEQRADNLSTIAHELQQYIDNLNNTLVDLEWLNRIETYVKWPWYVWLLIGL
EU186072_FCoV1_Black               LNFTAEIDQLEQRADNLTTIAHELQQYIDNLNKTLVDLEWLNRIETYVKWPWYVWLLIGL
AB088222_FCoV1_UCD1                LNLTAEIDQLEQRADNLTNIAHELQEYIDNLNKTLVDLEWLNRIETYVKWPWYVWLLIGL
                                   **:*****:*: *****: ***:**:******:*:***:****:****************

KP849472.1_CCoV1_Italy-23/03       VVIFCIPLLLFCCCSTGCCGCFGCIGSCCHSMCSRRQFESYEPIEKVHIH
AY307020.1_CCoV1_Elmo/02           VVIFCIPLLLFCCCSTGCCGCFGCIGSCCHSMCSRRQFESYEPIEKVHIH
AAB47503_FCoV1_KU-2                VVVFCIPLLLFCCLSTGFCGCFGCVGSCCHSLCSRRQFETYEPIEKVHIH
KX722530_FCoV1_Dutch_cat2          VVVFCIPLLLFCCLSTGCCGCFGCLGSCCNSFCSRRQFESYEPIEKVHIH
KX722529_FCoV1_Belgium_UG-FH8      VVVFCIPLLLFCCLSTGCCGCFGCLGSCCHSLCSRRQFENYEPIEKVHIH
KP143512_FCoV1_UK26M               VVVFCIPLLLFCCLSTGCCGCFGCLGSCCHSLCSRRQFESYEPIEKVHIH
MG893511_FCoV1_Germany_Felix       VVVFCIPLLLFCCLSTGCCGCFGCLGSCCHSLCSRRQFESYEPIEKVHIH
KY566209_FCoV1_China10             VVVFCIPLLLFCCLSTGCCGCFGCLGSCCNSLCSRRQFESYEPIEKVHIH
FJ938054_FCoV1_Utrecht_UU4         VVVFCIPLLLFCCLSTGCCGCFGCLGSCCHSLCSRRQFESYEPIEKVHVH
DQ848678_FCoV1_C1Je_Kitten_FIP     VLVFCIPSLLFCCLSTGCCGCFGCLGSCCHSLCSIRQFENYEPIEKVHIH
EU186072_FCoV1_Black               VIVFCIPLLLFCCLSTGCCGCFGCLGSCCHSLCSRRQFENYEPIEKVHIH
AB088222_FCoV1_UCD1                VVVFCIPLLLFCCLSTGCCGCFGCLGSCCHSLCSRRQFENYEPIEKVHIH
                                   *::**** ***** *** ******:****.*:** ****.********:*
```

**B**

**Summary of Amino Acid (AA) Sequence Identity and Similarity between CCoV1 and FCoV1**

| | **NTD** | **RBD** | **CTD (CCoV1)** | **CTD (FCoV1)** | **S1 (CCoV1)** | **S1 (FCoV1)** | **S2 (CCoV1)** | **S2 (FCoV1)** |
|---|---|---|---|---|---|---|---|---|
| Total No. of AA | 435 | 281 | 88 | 86 | 800 | 798 | 675 | 677 |
| No. of AA with Identity | 150 | 156 | 58 | 58 | 375 | 375 | 496 | 496 |
| No. of AA with Similarity | 253 | 227 | 71 | 71 | 567 | 567 | 603 | 603 |
| % Identity | 34.5% | 55.5% | 65.9% | 67.4% | 46.9% | 47.0% | 73.5% | 73.3% |
| % Similarity | **54.2%** | 80.8% | 80.7% | 82.6% | 70.9% | 71.0% | **89.3%** | **89.1%** |

#### S1 Table.
#### CD8$^+$ CTL and CD4$^+$ $T_H$ Epitopes on SCoV2 UF-RBD and UF2-RBD *

| U.S. Population / Race* | HLA-A/B Supertype / DRB1 Lineage | Allele / Allotype * | UF-RBD | UF2-RBD | |
|---|---|---|---|---|---|
| | | | No. of CTL / $T_H$ Epitope †‡ | No. of CTL / $T_H$ Epitope †‡ | N/C Extended Seq Epitope § |
| Both | A1 | A*0101 | 1 | 1 | 0 |
| Caucasian | A1 | A*3201 | 3 | 3 | 0 |
| Black | A1 | A*3001 | 3 | **5** | 1N+1C |
| Both | A2 | A*0201 | 1 | 1 | 0 |
| Caucasian | A2 | A*0205 | 2 | 2 | 0 |
| Black | A2 | A*6802 | 3 | 3 | 0 |
| Both | A3 | A*0301 | 4 | **8** | 3N+1C |
| Caucasian | A3 | A*1101 | 5 | **9** | 3N+1C |
| Black | A3 | A*7401 | 4 | **6** | 2N |
| | | **HLA-A Subtotal:** | 26 | 39 | 9N+3C |
| Both | B7 | B*0702 | 1 | **2** | 1C |
| Caucasian | B7 | B*5101 | 1 | 1 | 0 |
| Black | B7 | B*5301 | 2 | 2 | 0 |
| Caucasian | B44 | B*4402 | 1 | 1 | 0 |
| Caucasian | B44 | B*4001 | 2 | 2 | 0 |
| Black | B44 | B*4501 | 1 | **2** | 1N |
| Both | B44 | B*4403 | 1 | 1 | 0 |
| Caucasian | B27 | B*2705 | 3 | **4** | 1N |
| Both | B27 | B*1402 | 4 | **5** | 1N |
| Black | B27 | B*1503 | 5 | 5 | 0 |
| Caucasian | B27 | B*3901 | 2 | **3** | 1N |
| | | **HLA-B Subtotal:** | 23 | 28 | 4N+1C |
| | | **HLA-A+HLA-B Total:** | 49 | 67 | 13N+4C |
| Both | DRB1*04 | 0401 | 0 | 0 | 0 |
| Caucasian | DRB1*04 | 0404 | 0 | 0 | 0 |
| Caucasian | DRB1*04 | 0402 | 1 | 1 | 0 |
| Black | DRB1*04 | 0405 | 7 | 7 | 0 |
| Both | DRB1*15 | 1501 | 4 | 4 | 0 |
| Both | DRB1*07 | 0701 | 1 | 1 | 0 |
| Both | DRB1*03 | 0301 | 0 | 0 | 0 |
| Both | DRB1*13 | 1301 | 6 | 6 | 0 |
| Both | DRB1*13 | 1302 | 0 | 0 | 0 |
| Both | DRB1*01 | 0101 | 7 | 7 | 0 |
| Caucasian | DRB1*01 | 0103 | 0 | 0 | 0 |
| Both | DRB1*11 | 1101 | 13 | 13 | 0 |
| Both | DRB1*08 | 0801 | 6 | 6 | 0 |
| | | **HLA-DRB1 Total:** | 45 | 45 | 0 |

* Top 2-3 alleles or allotypes for U.S. Caucasian and Black populations according to the survey in the Allele Frequencies in Worldwide Populations database: http://www.allelefrequencies.net/hla6006a.asp Each individual has two HLA-A alleles, two HLA-B alleles, and two HLA-DRB1 alleles. The person will be protected the most if he/she has HLA-A and HLA-B alleles that can express allotypes which recognize many CD8$^+$ T-cell (cytotoxic T lymphocyte, CTL) epitopes.

† CD8$^+$ T-cell/CTL epitopes derived from NetMHCpan 4.1 server using 9mer peptide as the core. https://services.healthtech.dtu.dk/service.php?NetMHCpan-4.1
CD4$^+$ T-cell (T-helper, $T_H$) epitopes derived from NetMHCII 2.3 server using 15mer peptide as the core. https://services.healthtech.dtu.dk/service.php?NetMHCII-2.3

‡ These epitopes are not conserved epitopes and therefore are susceptible to mutation(s).

§ The number of T-cell epitope(s) present or overlapping with the extended 46-aa sequence (Seq) of the UF2-RBD at the amino-end (N) and with the extended 12-aa sequence at the carboxyl-end (C).
